# Bilateral spontaneous otoacoustic emissions show coupling between active oscillators in the two ears

**DOI:** 10.1101/484717

**Authors:** Y. Roongthumskul, D. Ó Maoiléidigh, A. J. Hudspeth

## Abstract

Spontaneous otoacoustic emissions (SOAEs) are weak sounds that emanate from the ears of tetrapods in the absence of acoustic stimulation. These emissions are an epiphenomenon of the inner ear's active process, which enhances the auditory system’s sensitivity to weak sounds, but their mechanism of production remains a matter of debate. To understand the relationship between SOAEs that we recorded simultaneously from the two ears of the tokay gecko, we developed a mathematical model of the eardrums as noisy nonlinear oscillators coupled by the air within a lizard’s mouth. We found that binaural emissions could be strongly correlated: some emissions occurred at the same frequency in both ears and were highly synchronized. Suppression of the emissions in one ear often changed the amplitude or shifted the frequency of emissions in the other. Decreasing the frequency of emissions from one ear by lowering its temperature usually reduced the frequency of the contralateral emissions. By according with the model, the results indicate that some SOAEs are generated bilaterally through acoustic coupling across the oral cavity. The model predicts that sound localization through the acoustic coupling between ears is influenced by the active processes of both ears.

## Introduction

One of the hallmarks of the auditory systems in all classes of tetrapods is the ability of inner ears to produce oscillations in the absence of external acoustic stimulation. Owing to the reverse transmission of sound through the middle ear, these oscillations elicit vibrations of the eardrum that are detected externally as weak sounds termed spontaneous otoacoustic emissions (SOAEs). From species to species, animal to animal, and ear to ear, these emissions vary in number, level, and frequency^1–6^.

SOAEs are sensitive to several types of manipulation. For example, the application of a pure-tone acoustic stimulus usually attenuates an emission and causes a shift in its frequency away from that of the stimulus frequency^4,7^. More rarely, some emissions are enhanced or shifted toward the stimulus^8^. Decreasing the body temperature evokes a decline in the frequency of emission^6,9^. Finally, altering the air pressure in the ear canal strongly affects SOAEs: extreme pressure changes typically attenuate the emissions. The associated shifts in SOAE frequency are complex and vary widely between species^10–12^. In addition to altering the acoustic impedance of the middle ear, changes in the ear-canal pressure might influence the intracochlear noise level or the active process underlying the emissions^11,13^.

Several studies report SOAEs emanating at nearly identical frequencies from both ears of an animal^14,15^. In mammals, the correlations between ears are speculated to arise from efferent control owing to the medial olivocochlear system. In amphibians, the strong binaural SOAEs are attributed to acoustic coupling between the ears^16^, for in most non-mammalian tetrapods the Eustachian tubes that connect the middle ears to the oral cavity are widely open. An external sound impinging on one tympanum can accordingly traverse the oral cavity and stimulate the contralateral ear^17–19^.

To evaluate the possibility that bilateral emissions at similar frequencies signify acoustic coupling between the two ears, we employed a model of nonlinear, active hair cells and inquired whether airborne coupling allows the simulated emissions to entrain one another. After modeling the effects of specific perturbations on emissions, we compared the model’s behavior with the results of binaural recordings from the tokay gecko, a convenient experimental preparation with robust SOAEs (Figure 1A,B).

**Figure 1.**
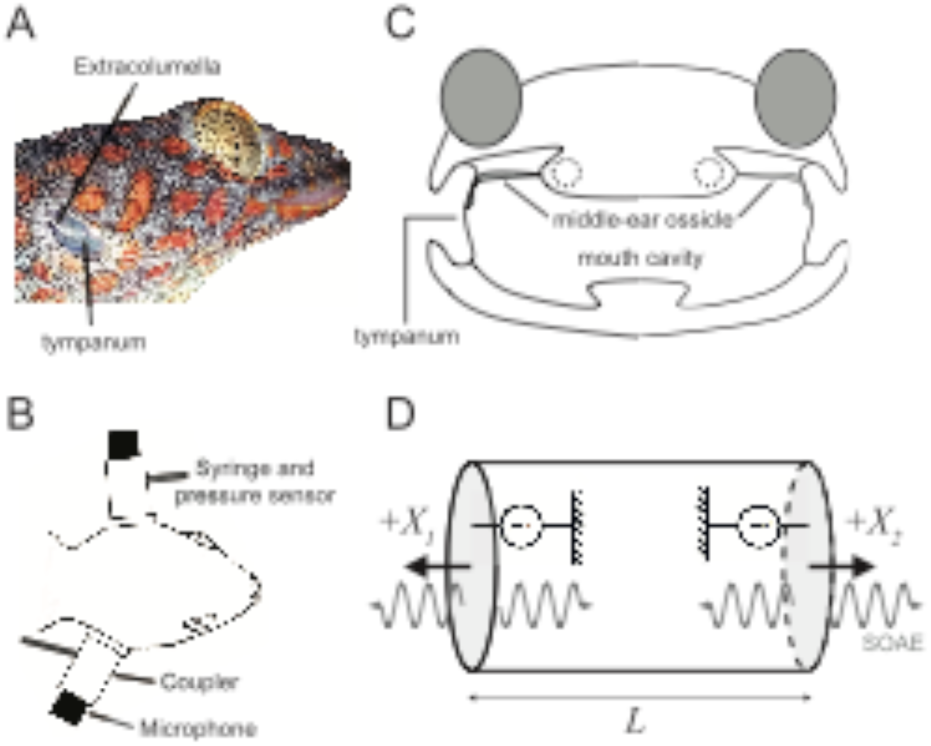
(A) The tympanum and the lateral articulation of the middle-ear ossicle, the extracolumella, are apparent through the tokay gecko's shallow ear canal. (B) A plastic coupler was sealed over each ear canal with vacuum grease. The coupler was attached to a sensitive microphone and connected by a plastic tube to a 5 mL syringe and pressure sensor. (C) A schematic drawing of a gecko’s head shows that the two eardrums are connected through the mouth cavity. Dashed circles indicate the locations of the inner ears. (D) A model of the ears as two coupled nonlinear oscillators. A closed cylinder representing the mouth cavity has both ends covered by eardrums, each directly connected to an active inner ear. The coordinate system is such that the positive direction corresponds to an outward motion of each eardrum. Standard parameter values used in the model are summarized in Table S1. Parameter values that differ from these standards are given in the relevant figure captions.

## Materials and Methods

### Numerical modeling of coupled nonlinear oscillators

We represent the mouth of a tokay gecko as a closed cylinder of length *L*, whose ends terminate in two eardrums of area *A*, each connected to an SOAE generator in the inner ear (Figure 1C,D). Because the mouth remained slightly open during experiments, the oral cavity might be described by two half-open cylinders closely apposed at their open ends. Near the fundamental frequency of the two-cylinder system, however, the eardrums move in antiphase, like the fundamental mode of a cylinder closed at both ends. Moreover, the harmonic frequencies of the head are much lower than that of a cylinder with a length equal to the head’s width^30^. We describe the head cavity as a cylinder closed at both ends, in which the effective length *L* is chosen so that the eardrums’ phase difference agrees with experimental observations (Fig. 2C).

**Figure 2.**
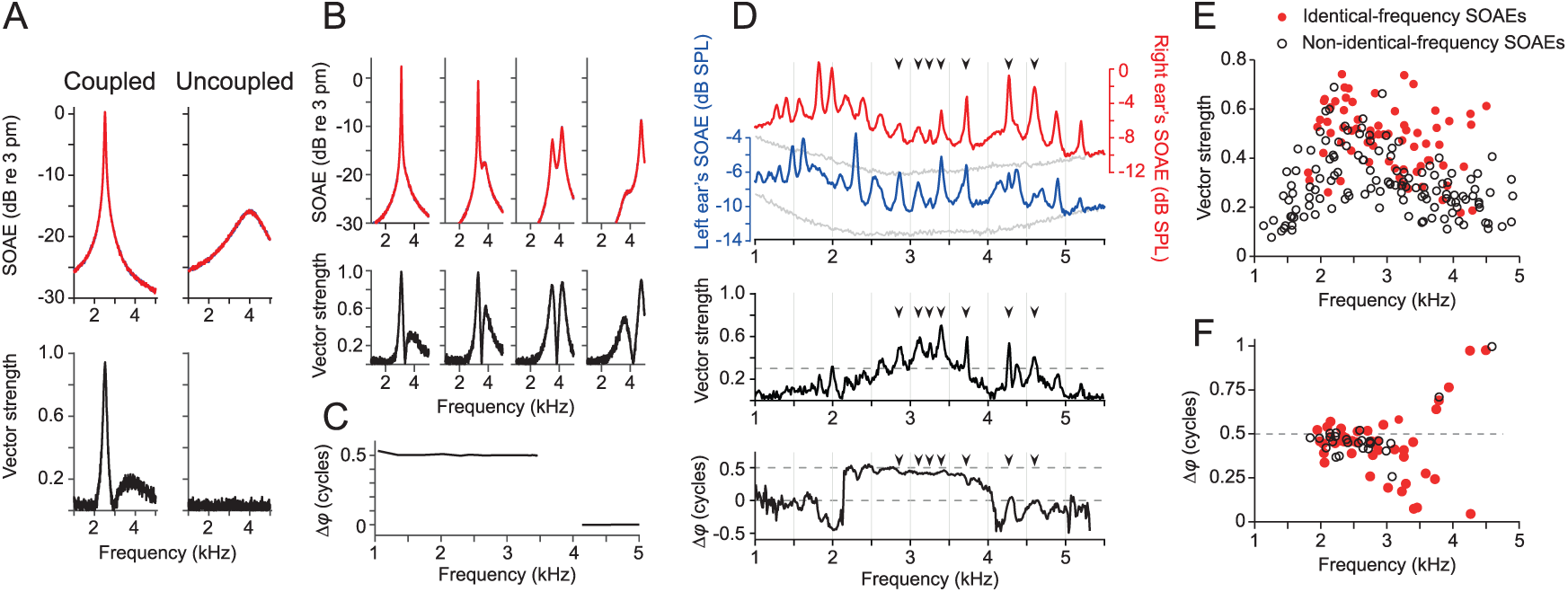
Identical-frequency spontaneous otoacoustic emissions (SOAEs), defined by a difference of less than 10 Hz in their center frequencies. (A) Coupling the oscillators to the head cavity reduces their oscillation frequency and increases their amplitude. The vector strength peaks at the oscillation frequency indicating synchronization at this frequency. The timescale parameter TS = 2. (B) The spectra and vector strengths of two identical oscillators are shown as their peak frequency is raised by reducing the timescale parameter: left to right, TS = 2, 1.5, 1.4, 1.3. When the peak frequencies are near the fundamental frequency of the head cavity (3.7 kHz), the acoustic coupling can create two spectral peaks. (C) The phase difference between two identical oscillators at their spectral peak is shown as a function of the peak frequency, which is set by adjusting the timescale parameter from 1.25 to 5.00. The oscillators move in antiphase below the fundamental frequency but in phase above the fundamental. Peaks do not appear close to the fundamental frequency. (D) The spectra of SOAEs recorded simultaneously from the left ear (blue) and right ear (red) display several identical-frequency SOAEs. The vector strength peaks at the frequencies of some SOAEs from both ears. The mean phase difference between binaural SOAEs peaks is close to 0.5 cycles below 3.5 kHz and is 0 cycles above 4 kHz. When the vector strength is small (<0.3), the phase difference is not constant and fluctuates considerably around the mean. Black arrowheads label the identical-frequency SOAEs with vector strength exceeding 0.3. The noise floors of the spectra (gray lines) were obtained from the same animal after euthanasia. (E) Most identical-frequency SOAEs (red dots) possess vector strengths exceeding 0.3. The vector strength sometimes exceeds 0.3 for non-identical-frequency peaks (black circles). (F) The phase difference between most identical-frequency SOAEs whose vector strength exceeds 0.4 is close to 0.5 cycles when the frequency is less than 3 kHz and is 0 cycles when the frequency exceeds 4 kHz. The data in panels E and F were obtained from 20 geckos.

The equation of motion of the nonlinear oscillator used to describe an eardrum’s displacement^13^ is

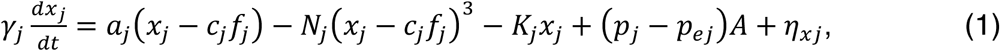

in which *x*_*j*_ denotes the displacement of the *j*^*th*^ oscillator (*j* = 1, 2). The oscillator is driven by the difference between the external pressure denoted by *p*_*ej*_ and the internal pressure *p*_*j*_ representing the pressure at the location of the *j*^*th*^ eardrum: *p*_1_ = *p*(0) and *p*_2_ = *p*(*L*). The first three terms on the right-hand side of the equation represent dynamics driven by active hair-bundle motility. The parameters *a*_*j*_, *b*_*j*_, and *K*_*j*_ are stifnesses, *c*_*j*_ is a compliance, *γ*_*j*_ is a damping coefficient, and *N*_*j*_ controls the strength of the system’s nonlinearity. *f*_*j*_ is an active force whose dynamics are given by

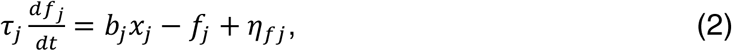

in which *τ*_*j*_ represents the timescale of the active process. The Gaussian white noise terms *η*_*x*_ and *η*_*f*_ satisfy

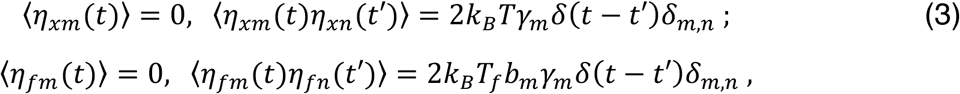

in which *k*_*B*_ is the Boltzmann constant. For an active system, the effective temperature *T*_*f*_ may differ from the thermodynamic temperature *T*^22^.

The motion of the air inside the cylinder is described in one dimension by its axial particle velocity *u*(*x, t*) and pressure *p*(*x, t*) with respect to a reference state of zero velocity, pressure *P*_r_ = 101 kPa, and temperature *T* = 25 °C. The air dynamics is described by the noisy acoustic telegraph equations

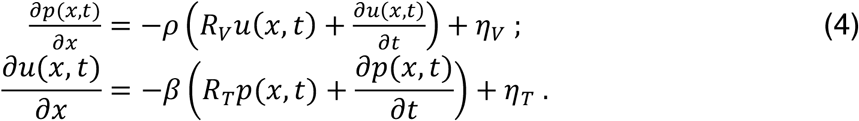

*R*_*V*_ and *R*_*T*_ are the viscous damping rate and the thermal damping rate, respectively. *ρ* is the air’s reference density and *β* is its adiabatic compressibility. The boundary conditions restrict the velocities of particles at both ends of the cylinder to equal those of the eardrums. The noise terms *η*_*V*_ and *η*_*T*_ satisfy

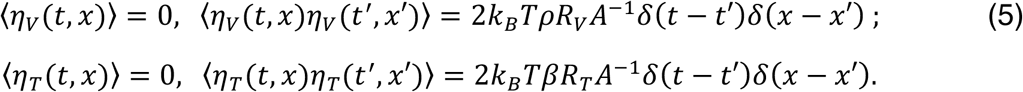

Numerical simulations of Equations 1-5 were performed by the forward Euler-Maruyama method (Equations 1 and 2) and through finite differences (Equation 4). To decrease the computational time, the small noise terms in Equation 4 were set to zero but are shown here for completeness. The solutions described the displacement of the eardrums *x*_*j*_. The emission frequency was identical to that of the oscillator, and the emission level was defined to be the displacement of the oscillator with respect to the reference level of 3 pm, a displacement chosen to match the velocity of the eardrum at 1 kHz and 0 dB SPL^17,31^. The steady-state time traces generated by the model were analyzed in the same fashion as the experimental time traces.

Other sources of dissipation in the actual system, such as the intricate anatomy of the mouth cavity and the presence of oral fluids, were accommodated through the use of damping rates larger than those corresponding to an unobstructed cylindrical cavity. More complex results could also arise from the interactions between SOAEs within an individual ear.

To change the frequency of an oscillator without changing its oscillation amplitude significantly, we used a dimensionless timescale parameter *TS* to rescale the oscillator’s time in Equations 1 and 2. Conversely, to change the amplitude with little change in the frequency, we employed the dimensionless parameter *DS* to rescale the oscillators’ displacement. These changes were achieved by rescaling the oscillators’ parameters as *a*_*j*_/*TS*^2^, *b*_*j*_/*TS*^2^, *K*_*j*_/*TS*^2^, *c*_*j*_*TS*^2^, *γ*_*j*_/*TS*, *N*_*j*_/(*TS*^2^*DS*^2^), and *τ*_*j*_*TS*. The values of all parameters are summarized in Table S1.

### Structure of the gecko's auditory system

The external ear of the tokay gecko consists of a shallow ear canal terminating in a conical tympanum whose tension is maintained by the single middle-ear ossicle, the columella, which attaches to the eardrum through a process termed the extracolumella (Figure 1A). Each internal ear includes a complex sensory epithelium, the basilar papilla, about 2 mm in length. The basal 700 μm of the organ is devoted to the frequency range 150-1000 Hz and includes some 200 hair cells^32^. The more apical 1300 μm of the basilar membrane, which represents frequencies from 1 kHz to more than 5 kHz^33^, supports two strips of hair cells. Beneath a narrow, continuous tectorial membrane lie 1000-1300 hair cells in rows of six to eight cells abreast; under 170 distinct sallets there are 800-900 hair cells in rows of five to seven cells abreast^32,34^.

### Measurement and manipulation of SOAEs

All experimental protocols were approved by the Animal Care and Use Committee of the Rockefeller University. Young adult tokay geckos (*Gekko gecko*) of both sexes were anesthetized with an intraperitoneal injection of 20 mg/kg sodium pentobarbital (Nembutal, Akorn Pharmaceuticals). The body temperature of the animal was maintained by a heating pad at 28.5 °C; the corresponding oral temperature was 24-25 °C. For all experiments, the distance between the tips of the upper and lower jaws was maintained near 10 mm.

Measurements of SOAEs were performed inside a sound-attenuation chamber with two microphones (ER-10B+, Etymotic Research), each connected to a 20 mm-long plastic coupler. The other end of each coupler was sealed over an animal's outer ear with vacuum grease (DC-976, Dow Corning). The air pressure inside each coupler was manually manipulated using a 5 mL syringe connected to the coupler by a plastic tube. The pressure level was monitored with a digital manometer (475-1-FM, Dwyer Instruments).

To reduce the temperature of an inner ear, we used a probe consisting of a 1.8 mm-diameter copper wire with one end attached to the cool surface of a Peltier cell (CP08, Laird Technologies). The other end of the wire was inserted into the mouth cavity and gently pressed against the mucosa overlying the ventral surface of the periotic bone of the left inner ear. The local temperature in the vicinity of each inner ear was monitored with a thermocouple placed against the mucosa abutting the periotic bone near the Eustachian tube.

### Calculation of SOAE spectra and peak detection

The pressure signal obtained from each ear was acquired in non-overlapping 100 ms segments at a sampling interval of 20 µs from 100 s recordings. The maximal pressure levels in each window and their standard deviations were calculated. To exclude extreme pressure variations owing to the animal’s respiratory movements, any sample whose peak pressure exceeded thrice the standard deviation was excluded from further analysis. A finite-time Fourier transform with a Hanning window was calculated for each of the remaining sections. The SOAE amplitude spectrum was obtained by averaging the magnitudes of the spectra.

Detection of SOAE peaks was performed on a smoothed emission spectrum with a five-point moving average. A peak was defined as a point in the spectrum whose magnitude exceeded those of the adjacent troughs by more than 45 nPa, which was equivalent to a peak of 0.2 dB above the noise floor at -20 dB SPL. This criterion was chosen such that the algorithm could detect a small identical-frequency SOAE superimposed upon another peak. The amplitudes of the emission spectra before averaging at the frequency corresponding to the peak had to differ statistically from those at both adjacent troughs by a Student’s *t*-test with a criterion of *p* < 0.2.

The phase difference of the binaural pressure signals was extracted from the finite-time Fourier transforms of each pair of non-overlapping 100 ms windows. The argument of a complex number *Z* is defined as 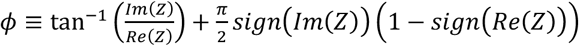. For each frequency component, the phase difference ∆*ϕ_j_*(*f*) between the complex Fourier components from both ears was given by the argument of the ratio 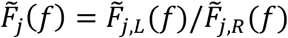, in which 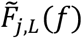 and 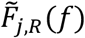 denote the complex Fourier component of the *j*^*th*^ segment of the signal from the left and right ear at frequency *f*. The phase difference of the mean ∆*ϕ*(*f*) was then obtained from the argument of the quantity 〈*e*^*i∆ϕ*_*j*_(*f*)^〉, in which 〈… 〉 denotes the ensemble average over all segments.

The vector strength was calculated as 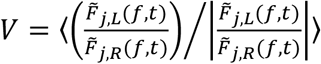. For perfectly phase-locked signals, the phase difference is time-invariant, and the vector strength becomes unity. If two emissions are independent the phase difference varies considerably with time and the vector strength is zero. Because the variance of the phase is small when the vector strength is low, the phase difference of the mean 〈*e*^*i∆ϕ*_*j*_(*f*)^〉 is meaningful only when the vector strength is large.

## Results

### A model of acoustically coupled noisy nonlinear oscillators

The detection of sound by tetrapods relies on the active process of hair cells, whose mechanosensitive hair bundles not only respond to deflections evoked by sound energy, but also enhance their oscillations through active movements^20,21^. Each hair bundle acts as a nonlinear oscillator whose dynamics is determined by its operating point^13,22,23^; over a specific range of operating points, the bundle exhibits enhanced sensitivity and frequency selectivity^24^. Under appropriate conditions an unstimulated bundle can exhibit spontaneous oscillations that might underlie SOAEs^25^.

To describe the synchronization of binaural emissions, we developed a mathematical model of two acoustically coupled nonlinear oscillators. In this model, each eardrum was driven by a noisy nonlinear oscillator based on active hair-bundle dynamics^13,24^. We assumed that the vibrations of an eardrum created not only ipsilateral SOAEs but also sound waves that traversed the mouth and caused a pressure difference across the contralateral tympanum; this provided acoustic coupling between the eardrums (Figure 1C, D). Propagation of the sound wave was governed by the damped-acoustic-telegraph equation^26^. The ensuing displacements of each eardrum were then used to calculate the corresponding SOAEs. For simplicity, we considered the phase-locking behavior of only one pair of binaural oscillators.

### Binaural correlation of SOAE spectra

Our modeling indicated that coupling of identical, active oscillators to the air-filled oral cavity and to each other increased the peak oscillation amplitudes and reduced the peak frequencies and bandwidths (Figure 2A). To quantify the degree of synchronization between the oscillators, we employed the vector strength, a metric for which a value of zero implies no phase-locking and a value of one implies perfect phase-locking. The maximal vector strength occurred at the oscillation frequency, indicating synchronization. The vector strength also exhibited a broad peak near the fundamental frequency of the head cavity, 3.7 kHz in the model, despite no evidence for emission peaks at that frequency. In contrast, uncoupled oscillators with identical oscillation frequencies were not synchronized. These results implied that ears should be considered strongly coupled at a particular frequency only if both ears exhibited emission peaks with high vector strength at that frequency. In this work, we defined synchronized emissions as those with identical frequencies and a vector strength exceeding 0.3.

We controlled the peak frequencies of both oscillators by adjusting a timescale parameter. Owing to the acoustic coupling, the emission spectra were bimodal when the peak frequency was close to the fundamental frequency of the oral cavity (Figure 2B). For oscillation frequencies below the fundamental frequency, the motions of the two eardrums were out-of-phase: as one eardrum moved toward the midline, the other moved away (Figure 2C). At higher frequencies, the eardrums moved in phase, a motion consistent with the second harmonic of the oral cavity.

The SOAE spectra recorded from a gecko’s ear featured several peaks at frequencies ranging from 0.5 kHz to 5.5 kHz (Figure S1). Comparison of the binaural spectra revealed that some emission peaks occurred at identical frequencies: their center frequencies differed by less than 10 Hz, the frequency resolution of the finite-time Fourier transform (Figures 2D and S2). The cross-correlation coefficient between the binaural spectra most often approached unity between 2 kHz and 4 kHz, indicating spectral similarity (Figure S3).

Investigation of the temporal correlations between the SOAEs of the two ears revealed that binaural emissions could exhibit a high degree of phase-locking over a broad frequency range. The vector strength displayed several peaks at frequencies corresponding to those of the emissions from both ears (Figure 2D). Peaks of high vector strength were associated primarily with identical-frequency SOAEs. According to our model, this observation signified strong coupling between the ears.

Analysis of the emissions recorded from 20 geckos revealed that identical-frequency peaks occurred only between 1.5 kHz and 4.5 kHz (Figure 2E). Identical SOAE frequencies were not always associated with phase-locking, however, for some possessed low vector strengths. In these instances, the ears coincidently oscillated at similar frequencies but were not strongly coupled. In contrast, we occasionally observed high vector strengths for frequencies at which an emission lacked a contralateral counterpart of the same frequency. Such a counterpart might have been excluded, however, by our conservative criterion for defining peaks as having identical frequencies.

The phase difference between most strongly coupled binaural emissions was 0.5 cycle for frequencies below 3 kHz and nil for frequencies above 4 kHz (Figure 2D, F). In agreement with the model, the frequency dependence of the phase difference can be explained by a fundamental frequency of the head cavity of 3-4 kHz.

### Pressure dependence of SOAEs

Our model suggested that the emissions from one oscillator would change upon suppression of the identical contralateral oscillator by the imposition of an external static pressure (Figure 3A). As the static force load shifted the contralateral oscillator into the quiescent region of the state space, its spontaneous oscillations declined^13^. This treatment often attenuated the emission level of the other oscillator. Depending on the oscillation frequency, the emission peaks of the unsuppressed oscillator increased or decreased in amplitude and frequency. Below about 5 kHz, the emission peaks shifted toward the fundamental frequency of 3.7 kHz when the contralateral oscillator was suppressed.

**Figure 3.**
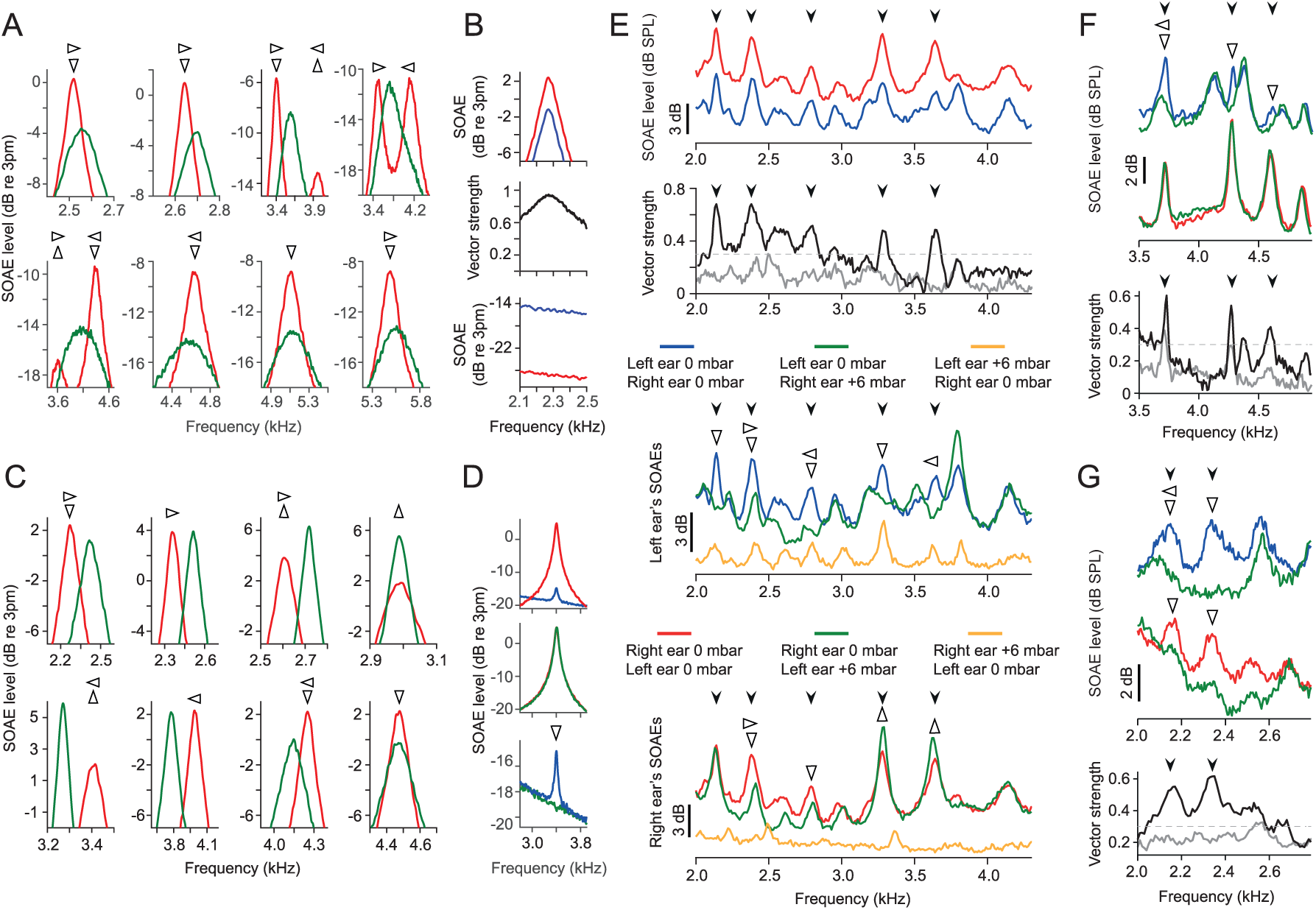
Effects of ear-canal pressure on SOAEs. An ear or oscillator was suppressed by applying an external pressure of +600 Pa. Changes in the emission level are identified by upright and inverted open arrowheads and frequency shifts are marked by sideways open arrowheads. (A) The spectrum of an active oscillator (red) changes when its identical contralateral counterpart is suppressed (green). As the oscillators’ peak frequency rises, the effects of suppression change. First row, left to right: TS = 2, 1.9, 1.45, 1.413. Second row, left to right: TS = 1.37, 1.32, 1.25, 1.2. (B) (Top, middle) Spontaneous oscillations of an active oscillator (red) entrain a passive oscillator (blue). (Bottom) Applying an external static pressure to the active oscillator eliminates oscillations of both oscillators. Passive oscillator: TS = 20. Active oscillator: TS = 2 and K = 0.7K_H_. (C) An active oscillator (red) drives a passive oscillator. In response to suppression of the passive oscillator, the change in the active oscillator’s peak depends on its peak frequency (green). First row, left to right: TS = 2, 1.95, 1.8, 1.63. Second row, left to right: TS = 1.5, 1.355, 1.3, 1.255. (D) (Top) The passive oscillator’s effect on the active oscillator is reduced by lowering its displacement scale parameter DS. (Middle) Suppression of the passive oscillator has little effect on the active oscillator. (Bottom) Suppression of the active oscillator eliminates emissions from both oscillators. Passive oscillator: TS = 20, DS = 0.05. Active oscillator: TS = 1.45 and K = 0.7K_H_. (E) (Top) SOAEs were recorded from the left ear (blue line) and the right ear (red line) when both ear canals were at atmospheric pressure. (Second) The vector strengths of most identical-frequency SOAEs are relatively large (black line) and diminish when the emissions from either ear are suppressed (gray line). (Third) SOAE spectra were recorded from the left ear as the pressure was raised to +600 Pa in the right ear canal (green line) or in the left ear canal (orange line). (Bottom) SOAE spectra were recorded from the right ear as the left ear-canal pressure was at +600 Pa (green line), and the right ear-canal pressure was at +600 Pa (orange line). (F, G) (Top) For two animals, the top panels illustrate emission spectra recorded from the left ear when the right ear canal was at atmospheric pressure (blue line) or at +600 Pa (top green line). (Middle) SOAEs from the right ear were recorded when the left ear canal was at atmospheric pressure (red line) or at +600 Pa (bottom green line). (Bottom) The vector strengths before (black line) and after (gray line) the external pressure was altered. In panels E-G, black arrowheads indicate identical-frequency SOAEs whose vector strength exceeds 0.3.

Ears might be considered active within the frequency ranges corresponding to emission peaks and behave as if they were passive at other frequencies. We modeled one oscillator as active and the other as passive but compliant by increasing its timescale parameter. In this case, the active oscillator drove emissions emanating from both ends of the cylindrical cavity. Suppression of the active oscillator caused the emissions to vanish from both ears (Figure 3B). Suppression of the passive oscillator shifted the frequency and modulated the amplitude of emissions from the active oscillator (Figure 3C). At some frequencies and when the displacement of the passive oscillator was sufficiently small, however, suppression of the passive oscillator had little effect on the active one (Figure 3D).

To experimentally investigate the effects of contralateral suppression of SOAEs, we used a plastic syringe connected to the microphone coupler to control the static pressure within each ear canal (Figure 1B). All emissions were attenuated with an increase or decrease in the ear-canal pressure (Figure S4). Suppression was typically achieved as the pressure exceeded +600 Pa to +800 Pa, at which level the emission spectra approached the noise floor (Figure 3E-G).

Suppression or substantial attenuation of emissions altered the SOAE spectrum recorded from the ear contralateral to the pressure change (Figures 3 and S5). The frequency and amplitude of an emission rose, fell, or were invariant upon suppression of the contralateral ear. Consistent with the emissions generated in the model of two active oscillators (Figure 3A), we occasionally encountered SOAE peaks that were detectable only in the presence of the emissions from both ears and vanished upon suppression of the emission from either (Figure 3G). These SOAEs typically exhibited a very high vector strength.

By comparing the emission amplitudes obtained before and during suppression of the contralateral SOAEs, we tested the statistical significance of the changes in emission spectra. The SOAE level exhibited significant changes (*p* < 0.001 by Student’s *t*-test) in response to suppression of the contralateral emissions (Figure S6).

### Effect of frequency detuning on SOAEs

To determine more concretely whether some emissions arose from the interaction of two active oscillators, we increased the peak frequency of the left oscillator in our model while that of the right stayed constant. The peak amplitude of and entrainment between the two oscillators rose as their frequency detuning declined (Figure 4A). As a clear indication of synchronization between the active oscillators, the peak frequency of the right oscillator shifted toward that of the left (Figure 4B). Over a limited range of small frequency detuning, synchronization of active oscillators was evidenced by their peak frequencies becoming identical and vector strength achieving a maximum (Figure 4C). When the oscillators were sufficiently well synchronized their peak frequencies rose at the same rate as functions of the timescale parameter. The peak frequency of the right oscillator then returned to its unperturbed value as the left oscillator’s influence fell with increased detuning.

**Figure 4.**
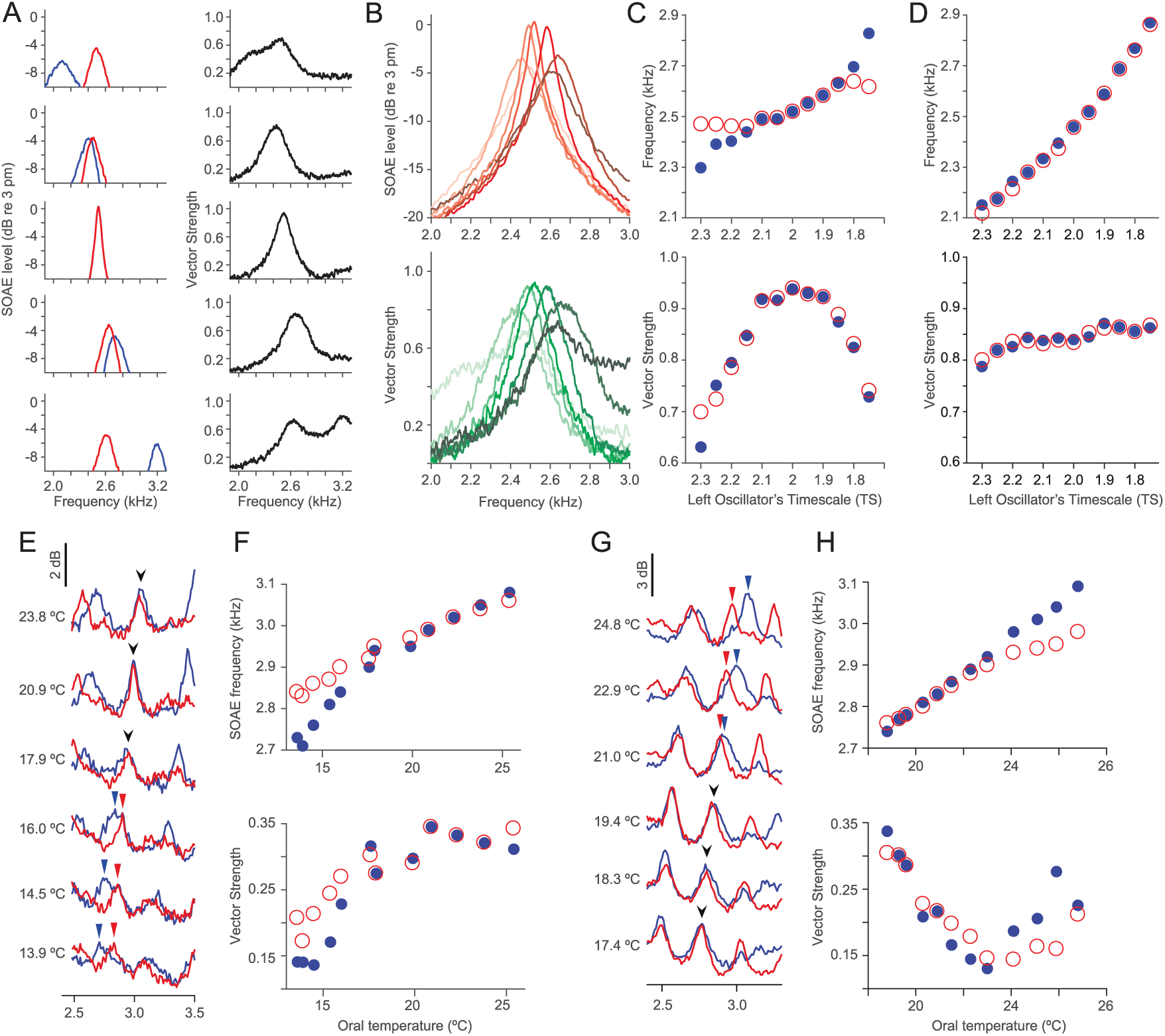
Synchronization of active-oscillator emissions. (A) The amplitudes and vector strengths of emissions from two acoustically coupled active oscillators are shown as a function of the emission frequency for different values of the left oscillator’s timescale parameter. Right oscillator (red): TS = 2. Left oscillator (blue), top to bottom: TS = 2.5, 2.2, 2, 1.8, 1.6. (B) The spectral amplitude of the right unperturbed oscillator and the vector strength are shown as functions of frequency. Light to dark: TS = 2.5, 2.2, 2.1, 2.0, 1.9, 1.8, 1.6. (C) The peak emission frequency and vector strength at the peak frequency are shown as functions of the left oscillator’s timescale parameter (left oscillator, blue dots; right oscillator, red circles). (D) The peak emission frequency and vector strength at the peaks are shown as functions of the active left oscillator’s timescale parameter when the right oscillator is passive (left oscillator, blue dots; right oscillator, red circles). Right oscillator: TS = 20. (E) SOAEs were recorded simultaneously from the left ear (blue lines) and right ear (red lines) at various temperatures of the left ear. As the temperature was reduced, a pair of identical-frequency SOAEs at 3 kHz (black arrowheads) separated into two non-overlapping emission peaks (blue arrowheads for left ear and red arrowheads for right ear). (F) The frequencies and vector strengths of SOAE peaks from the left ear (blue dots) and the right ear (red circles) are shown as functions of the left ear’s temperature. (G) In another animal, cooling caused two distinct SOAE peaks (blue and red arrowheads) to converge into a pair of identical-frequency SOAEs (black arrowheads). (H) The frequencies of the SOAE peaks converged and the vector strengths increased as the temperature declined.

In contrast, when an active left oscillator drove a passive right oscillator, their peak frequencies remained similar at all parameter values and the vector strength did not exhibit a maximum (Figure 4D). The intrinsic rate at which the left oscillator’s frequency rose was relatively insensitive to the value of the timescale parameter and was on average greater than when both oscillators were active (Figure 4C).

### Temperature dependence of SOAEs

Previous studies revealed that the frequency of SOAEs from amphibians and reptiles depends strongly on body temperature^4,9,27^. We confirmed experimentally that all SOAE peaks displayed negative shifts in frequency upon a reduction in temperature (Figure S7). The magnitude of the shift increased monotonically as a function of the emission frequency and could be described empirically by an exponential function^27^. We took advantage of this feature by reducing the local temperature of one inner ear with a cooling probe inserted into the oral cavity and pressed against the mucosa overlying the temporal bone. As the left inner ear was cooled, the right ear’s temperature also decreased owing to thermal conduction, but the magnitude of the contralateral effect was significantly smaller (Figure S8). We were thus able to impose asymmetrical cooling on the two inner ears and to investigate the resulting effects on binaural emissions.

Asymmetrical reductions in temperature showed that a pair of synchronized emissions could separate into two weakly correlated emissions with distinct frequencies (Figure 4E). Over a moderate range of temperature changes at the left ear, these identical-frequency SOAEs underwent similar frequency shifts (Figure 4F). Below some critical temperature, however, the emissions changed at different rates with respect to the left ear’s temperature, which led to their dissociation into two distinct SOAEs. The separation of emissions was concomitant with a decrease in the vector strength. Reducing the temperature also allowed some uncorrelated SOAEs to converge (Figure 4G). As the temperature decreased the emissions first aligned in frequency before they became synchronized (Figure 4H). These observations accorded with the model's behavior when both oscillators were active and consequently implied that some emissions arose from the synchronization of two active ears.

Because emissions from both ears exhibited changes in frequency as one ear was cooled, we investigated the temperature dependence of both SOAE spectra in more detail. When the left ear was cooled, binaural SOAE spectra revealed that most emissions from that ear exhibited greater changes in frequency than did those from the right ear (Figure 5A). However, some highly phase-locked identical-frequency SOAEs observed in the left ear showed significantly smaller shifts than the neighboring peaks (Figure 5B). Some highly synchronized emission peaks from the right ear likewise underwent unusually large frequency shifts. The different temperature sensitivities of neighboring peaks occasionally led to their crossing or coalescence.

**Figure 5.**
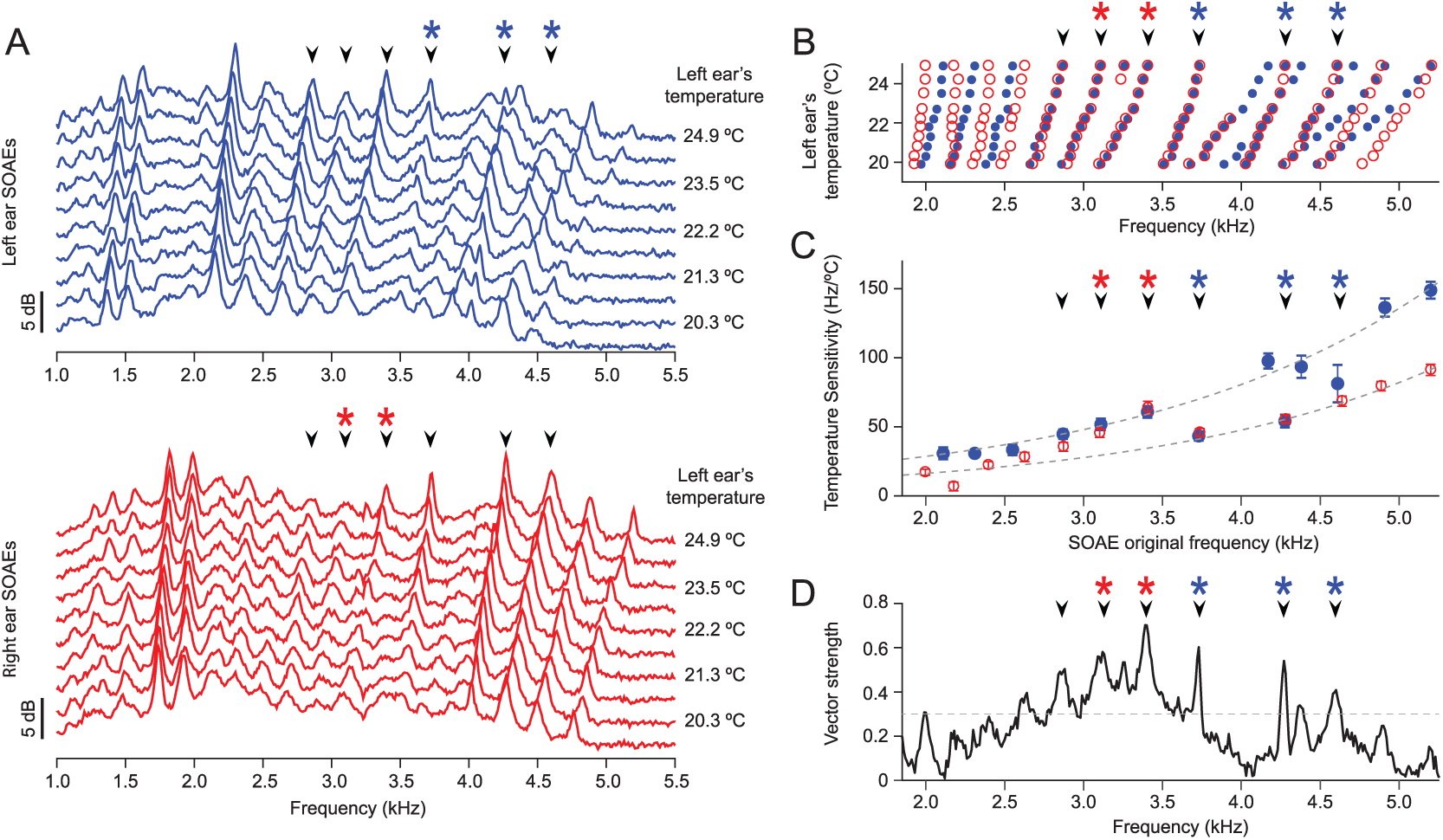
Asymmetric thermal manipulation of SOAEs. In all panels, black arrowheads label the identical-frequency SOAEs whose vector strength exceeded 0.3. (A) The spectra of SOAEs from the left ear (blue lines, top panel) and the right ear (red lines, bottom panel) were recorded as the left ear was cooled. The temperature adjacent to the left inner ear is indicated to the right of each plot. Data were obtained from the same animal as that for Figures 2D and 3F. (B) The center frequencies of all SOAE peaks from the left ear (blue dots) and right ear (red circles) showed systematic shifts to lower frequencies during cooling. (C) The center frequencies of SOAEs from the left ear (blue dots) and right ear (red circles) displayed distinct sensitivities to the temperature of the left ear. The temperature sensitivities are the inverse of the line slopes in panel B and define the peak frequencies as a function of the left ear’s temperature. Error bars indicate the 95 % confidence intervals of the fits. After exclusion of identical-frequency SOAEs with high vector strength, the intrinsic temperature sensitivities of each ear could be described by the exponential functions (gray dashed lines): *df*/*dT* = 9.933*e*^*f*_0_/1917^ (left ear) and *df*/*dT* = 5.303*e*^*f*_0_/1832^ (right ear). *f*_0_ denotes the original SOAE frequency at 24.9 °C. The temperature sensitivity of some identical-frequency SOAE peaks in the left ear (blue asterisks) deviated from the intrinsic exponential curve and equaled that of their counterparts in the right ear. Certain peaks in the right ear (red asterisks) exhibited deviant temperature sensitivities that accorded with those of their contralateral counterparts. (D) The vector strengths of SOAEs at 24.9 °C indicate that the peaks exhibiting deviant frequency responses were highly phase-locked.

Because the temperature dependence of each SOAE frequency was approximately linear (Figure 5B), the temperature sensitivity of each peak could be quantified as the rate of change of the peak frequency with respect to the left ear’s temperature. For weakly phase-locked peaks, the temperature sensitivity increased exponentially as a function of the original SOAE frequency at 24.9 °C (Figure 5C,D). The exponential increase with emission frequency defined the intrinsic temperature sensitivity of each ear. Because the left ear’s temperature was changed directly, the temperature sensitivities of weakly phase-locked emissions from that ear exceeded those of the right ear. In contrast, the temperature sensitivities of some highly synchronized peaks equaled those of the contralateral ear, a phenomenon that was observed for multiple geckos (Figure S9). The temperature sensitivities of highly synchronized emissions deviated from the intrinsic sensitivity curve of each ear.

To understand why emissions sometimes followed the temperature sensitivity of the ipsilateral ear and sometimes that of the contralateral ear, we implemented a model in which the left oscillator was weaker than the right (Figure 6A). Because the right oscillator dominated and entrained the left, the left oscillator’s emission frequency was insensitive to the value of its timescale parameter. When the right oscillator was then immobilized by increasing its stiffness substantially^13^, the left oscillator’s sensitivity to its timescale parameter was restored. In contrast, if the left oscillator was more robust than the right, immobilizing the right oscillator had little effect on the left oscillator’s sensitivity to changes in its parameter value (Figure 6B).

**Figure 6.**
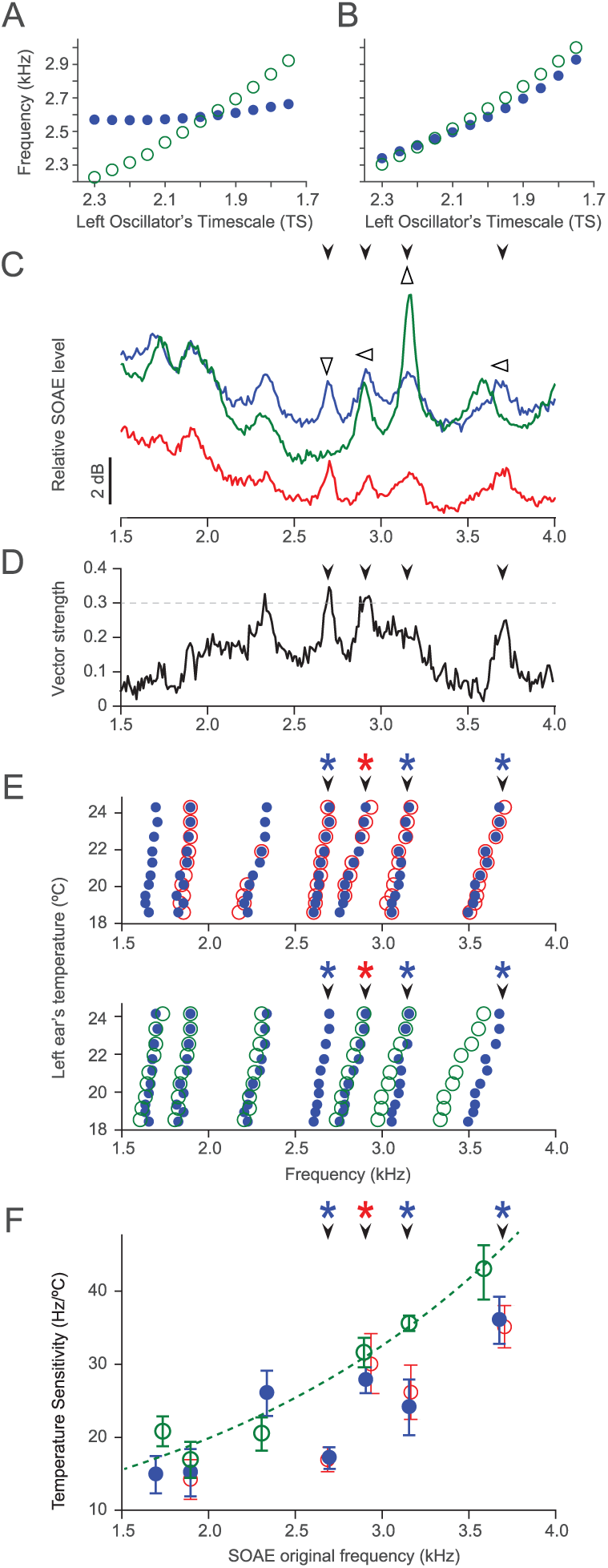
Influence of the contralateral ear on the parameter dependence of SOAEs. (A) Because the stronger right oscillator (DS = 2) dominates and entrains the left oscillator, the left oscillator’s emission frequency changes little as its timescale parameter is changed (blue dots). When the right oscillator is immobilized by raising its stiffness (K = 10K_H_), the left oscillator’s emission frequency changes at its intrinsic sensitivity to the left timescale parameter (green circles). (B) When the right oscillator is weaker than the left oscillator (DS = 2), the left oscillator’s sensitivity to its timescale parameter is almost independent of whether the right oscillator is immobilized (green circles, K = 10K_H_) or not (blue dots). (C) After recording of control SOAEs from the left ear (blue line) and right ear (red line), we suppressed the right ear’s emissions by immobilizing the columella. Some emissions from the left ear (green line) then displayed qualitative changes (open arrowheads). (D) A plot of vector strength reveals that not all identical-frequency SOAEs sensitive to contralateral suppression exhibited high vector strengths (>0.3). In panels C-F, black arrowheads label the identical-frequency SOAEs that are sensitive to contralateral suppression. (E) The frequencies of some SOAE peaks shown in panel C for the left ear (blue dots) and the right ear (red circles) displayed similar sensitivities to the temperature of the left ear. After immobilization of the right columella, some of the left ear’s emissions changed their dependence on temperature (green circles). (F) Before immobilization, the temperature sensitivity of a peak from the right ear deviated from the intrinsic sensitivity of that ear (red asterisk) and the left ear’s temperature sensitivities did not rise monotonically with the emission frequency. Upon immobilization of the right columella, the temperature sensitivities of the left ear’s emissions increased monotonically with the emission frequency (green circles). The left ear’s peaks with originally deviant temperature sensitivities either vanished or rose to agree with that ear’s intrinsic temperature sensitivity curve (blue asterisks).

To experimentally suppress emissions by increasing an eardrum's stiffness, we applied cyanoacrylate tissue adhesive at the base of the columella of the right ear, near its insertion onto the inner ear. As a result, some emissions vanished whereas others grew in magnitude and shifted in frequency (Figure 6C,D). These effects were similar to those obtained from increasing the ear-canal pressure.

Before suppression, the temperature sensitivity of the left ear's emissions did not rise monotonically with frequency (Figure 6E,F). When we cooled the left ear after immobilization of the right columella, however, the temperature sensitivity increased monotonically. For the left ear, the temperature sensitivities of peaks that were previously close to those of the right ear grew to match the temperature sensitivities of the weakly phase-locked emissions.

The model allowed us to interpret even these complex results. Two of the left ear’s peaks with temperature sensitivities close to those of the right ear were the result of a strong emission from the right ear dominating a weaker emission from the left. The emission that vanished upon immobilization of the right ear was created by the right ear driving the left, which did not oscillate spontaneously at that frequency. Finally, one emission peak resulted from the left ear driving the weaker right ear, for its temperature sensitivity was minimally affected by suppression of the right ear’s emission.

### Modeling the effects of interaural coupling on active sound localization

It is thought that acoustic coupling of the ears though the oral cavity allows a tokay gecko to localize sounds of wavelengths exceeding the animal's head size^28^. Because the detection of directionality has been explored for only high sound levels, at which the ear’s active process has little effect, it is unclear whether spontaneous activity can enhance the localization of weak sound sources^17^. To investigate this issue, we analyzed the response of the model to external stimuli. Both oscillators were stimulated with external sinusoidal pressure signals of the same frequency and magnitude. We assumed that the sound source was adjacent to one ear; to account for the gecko’s head size, we delayed the signal to the contralateral ear by 50 μs.

Even when the oscillators were identical, their responses to an external signal, measured at the stimulus frequency, depended on the sound source’s location (Figure 7A). When the source was adjacent to the left oscillator and the stimulus frequency lay below the oscillators’ emission frequency, the response of the left oscillator was smaller than that of the right oscillator. The converse was true for stimuli with frequencies above the emission frequency. The responses at various stimulus frequencies illustrated the ability of such a system to distinguish the direction of the stimulus over a broad range of frequencies (Figure 7B). The difference in the bilateral response was more pronounced when the oscillators were not identical (Figure 7C). Near the peak emission frequency of an oscillator, the size of the contralateral oscillator’s response depended on the location of sound source. Stimulation with a frequency far from an oscillator’s peak emission frequency evoked a minimal response when the source was placed next to the contralateral ear (Figure 7D). These results suggest that the acoustical coupling of active ears allows an animal to determine the location of weak high-frequency sounds.

**Figure 7.**
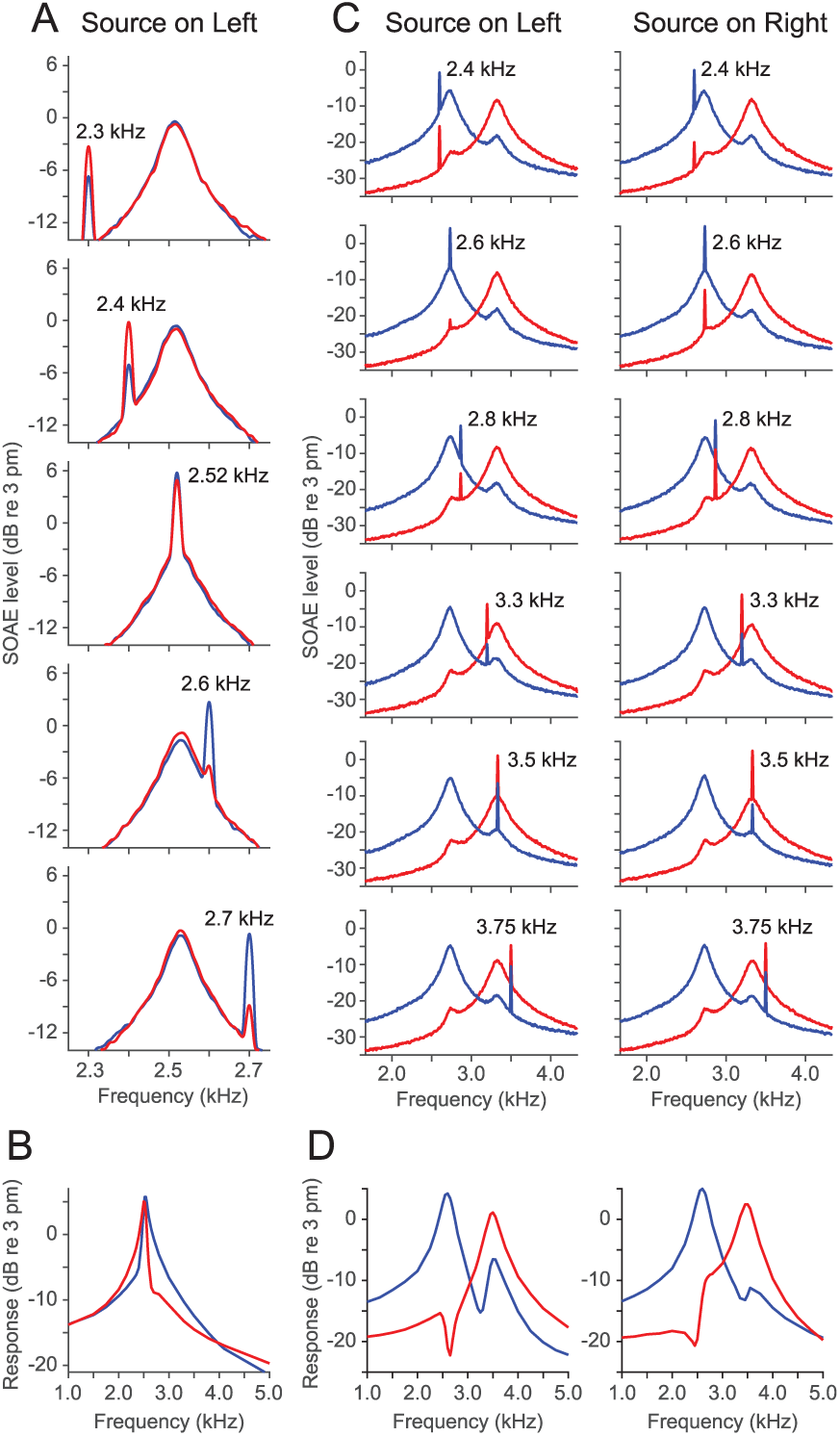
Sound source localization by acoustically coupled active oscillators. In the model system, a 20 dB SPL sound source is adjacent to one oscillator. To account for the travel time around the head to the other oscillator, the unattenuated signal at the contralateral oscillator is delayed by 50 μs. (A) The displacement spectra of two identical oscillators (TS = 2) are shown for different stimulus frequencies. The source is adjacent to the left oscillator, so the right oscillator’s stimulus is delayed. (B) Responses of identical oscillators (TS = 2) at the stimulus frequency are shown as functions of the stimulus frequency. (C) The displacement spectra of oscillators with distinct properties are shown when the source is adjacent to the left oscillator (TS = 2) or right oscillator (TS = 1.5). (D) The response at the stimulus frequency is shown when the source is adjacent to either the left oscillator (TS = 2) or the right oscillator (TS = 1.5). The responses of the left oscillator are shown in blue and those of the right oscillator in red.

## Discussion

We observed synchronization between the binaural SOAEs recorded simultaneously from the ears of the tokay gecko. Manipulations of one ear indicated that some SOAEs strongly influenced the emissions of similar frequencies from the contralateral ear. In some instances these effects could be explained only by the interaction of two active oscillators. These observations thus add considerable evidence for SOAEs being produced by the ear's active process.

Unlike mammalian cochleas, the auditory organs of most lizard species—including geckos—lack efferent innervation in the high-frequency region responsible for the generation of SOAEs. Because a model of the ears as acoustically coupled active oscillators agreed well with the experimental observations, the interaural coupling observed in this work likely occurred at the peripheral level and stemmed from acoustic coupling between the eardrums.

In agreement with the experimental observations, our model captured the acoustic synchronization of the oscillators and the frequency-dependent phase differences between ears. The model enabled us to interpret the complex alterations in emission upon suppression of the contralateral ear by static pressure and the change in emission frequency as the contralateral oscillator’s peak frequency was varied. Modeling also implied that the synchronization of emissions depended on the detuning in their peak frequencies and the emission strengths of both ears. The model predicted that increases in stiffness and raising or lowering the constant force created by static pressure suppresses spontaneous oscillations in the ear^13^. That these predictions have been confirmed experimentally for individual hair bundles^24,29^ and now at the whole-organ level provides compelling evidence that some SOAEs arise from the spontaneous oscillations of hair bundles.

Because the spontaneous activity of the two inner ears of a lizard can be synchronized, the active process of each ear affects the dynamics of both. Although internally coupled ears are thought to encode the location of high-intensity sound sources through the motions of two acoustically coupled eardrums, this effect has not been investigated for weak sounds^17–19^. Our mathematical model predicts that the activities of both ears dictate the sensitivity of each to weak sounds and consequently determine how each ear encodes the location of a sound source. The two ears of a gecko evidently function together as a single active system that is sensitive to the location of weak sound sources.

## Authors Contributions

Y.R. initiated the project, conducted physiological recordings, and wrote the manuscript; D.Ó M. carried out the theoretical analysis and modeling and wrote the manuscript; A.J.H. wrote the manuscript.

## Supplementary Material

**Figure S1.**
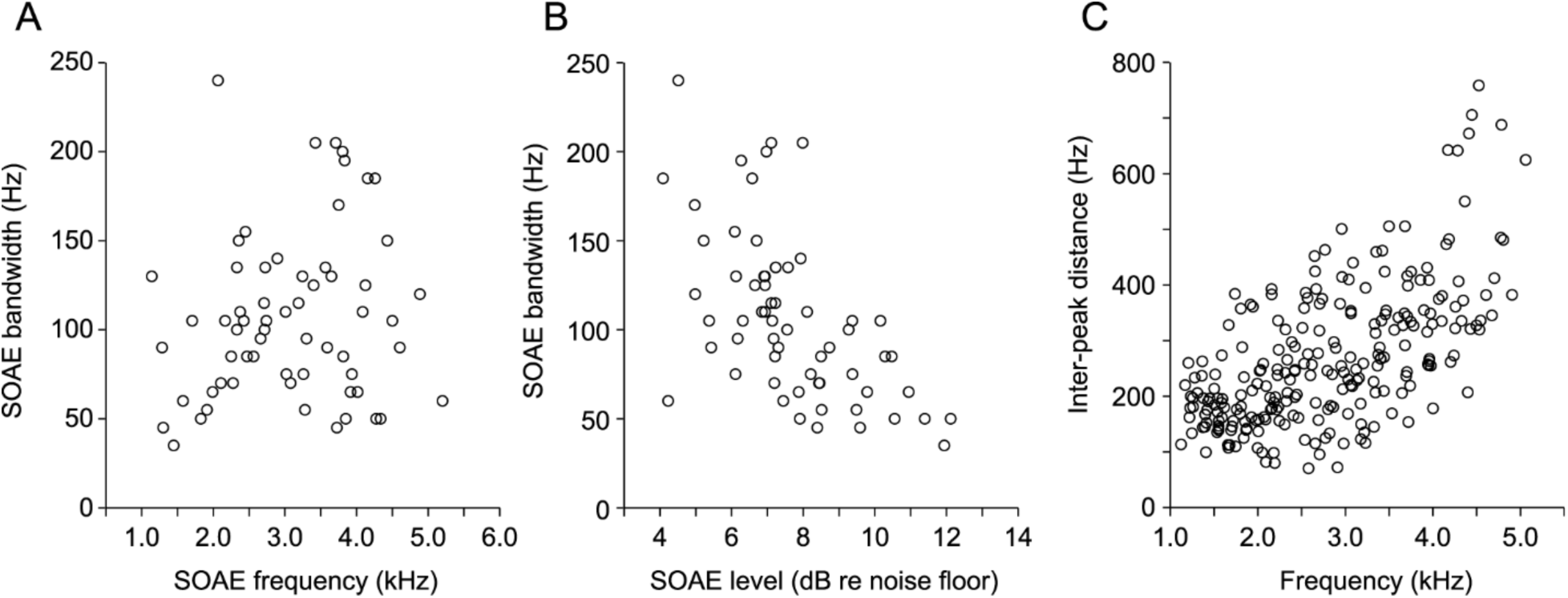
Characteristics of SOAEs from the tokay gecko. (A) The 3 dB bandwidth of each emission peak displays a weak dependence on SOAE frequency. (B) The bandwidth decreases as a function of SOAE level in decibels relative to the noise floor. Emissions with amplitudes less than 3 dB above the noise floor were excluded from the plots. (C) The distance between two adjacent SOAE peaks as a function of their average frequency. All SOAEs detected by means of the peak-detection algorithm were included in the analysis.

**Figure S2.**
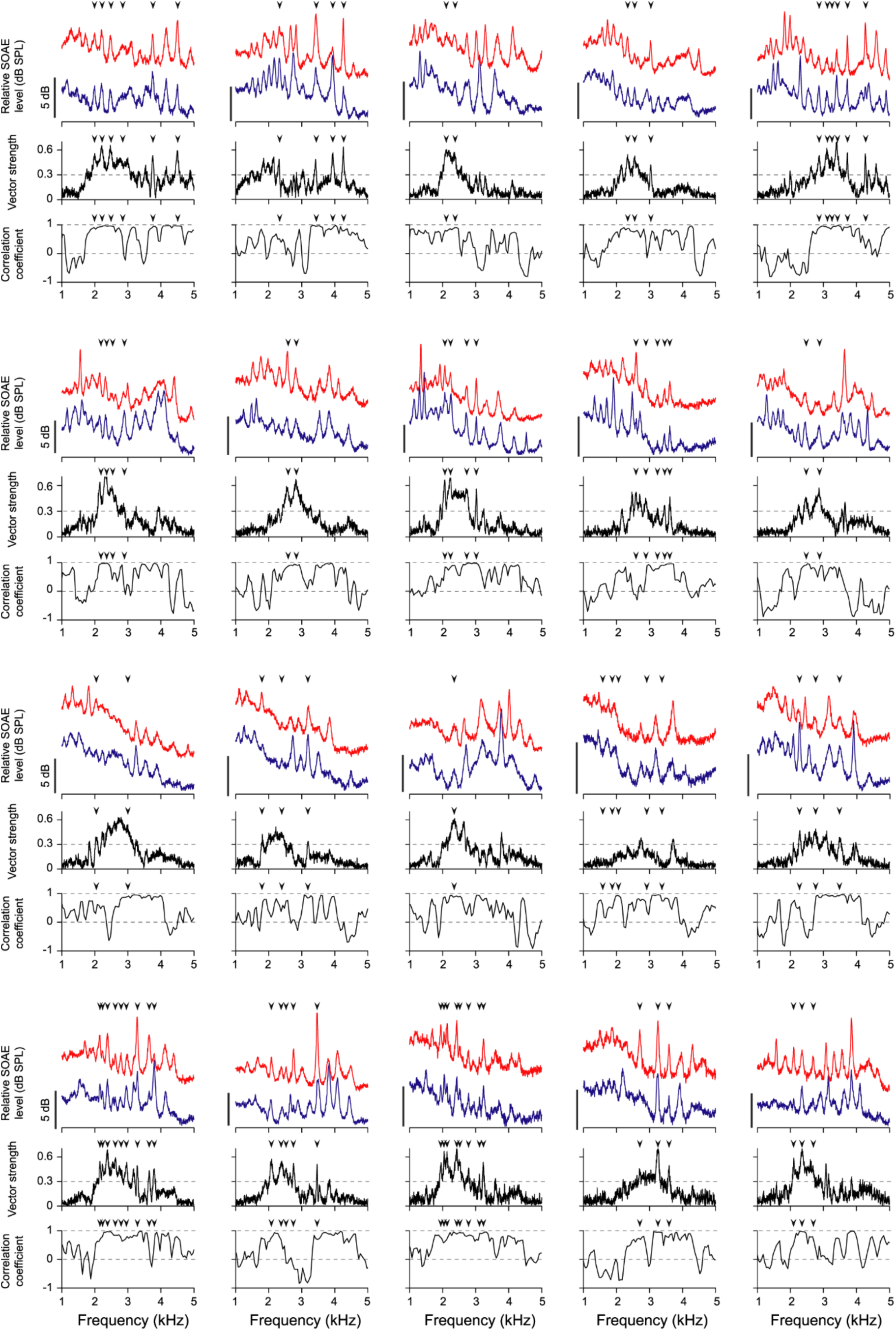
Binaural SOAE spectra recorded from 20 geckos. In each panel, the top traces illustrate the emission spectra recorded simultaneously from the left ear (blue line) and the right ear (red line). The binaural SOAE spectra are offset vertically to minimize overlap. The vector strengths of the phase differences between the ears are shown in the middle plots. A high degree of phase-locking is typically observed between 2 kHz and 3 kHz. The bottom plots show the cross-correlation coefficients calculated from the bilateral emission spectra. High spectral similarity and synchronization are observed within roughly the same frequency range. Black arrowheads indicate identical-frequency SOAEs whose vector strength exceeds 0.3.

### Calculation of the cross-correlation coefficient

To quantify the binaural spectral similarity, a cross-correlation coefficient was calculated from two 250 Hz windows, one from each spectrum over the same frequency range. Both windows were shifted together by 50 Hz increments to compute the coefficients at different frequencies.

We found that most pairs of identical-frequency SOAEs displayed large correlation coefficients, except those that differed in frequency from a neighboring monoaural peak by significantly less than our analysis window width. The average cross-correlation coefficients indicated high spectral similarity within the mid-frequency range of the spectra (Figure S3).

**Figure S3.**
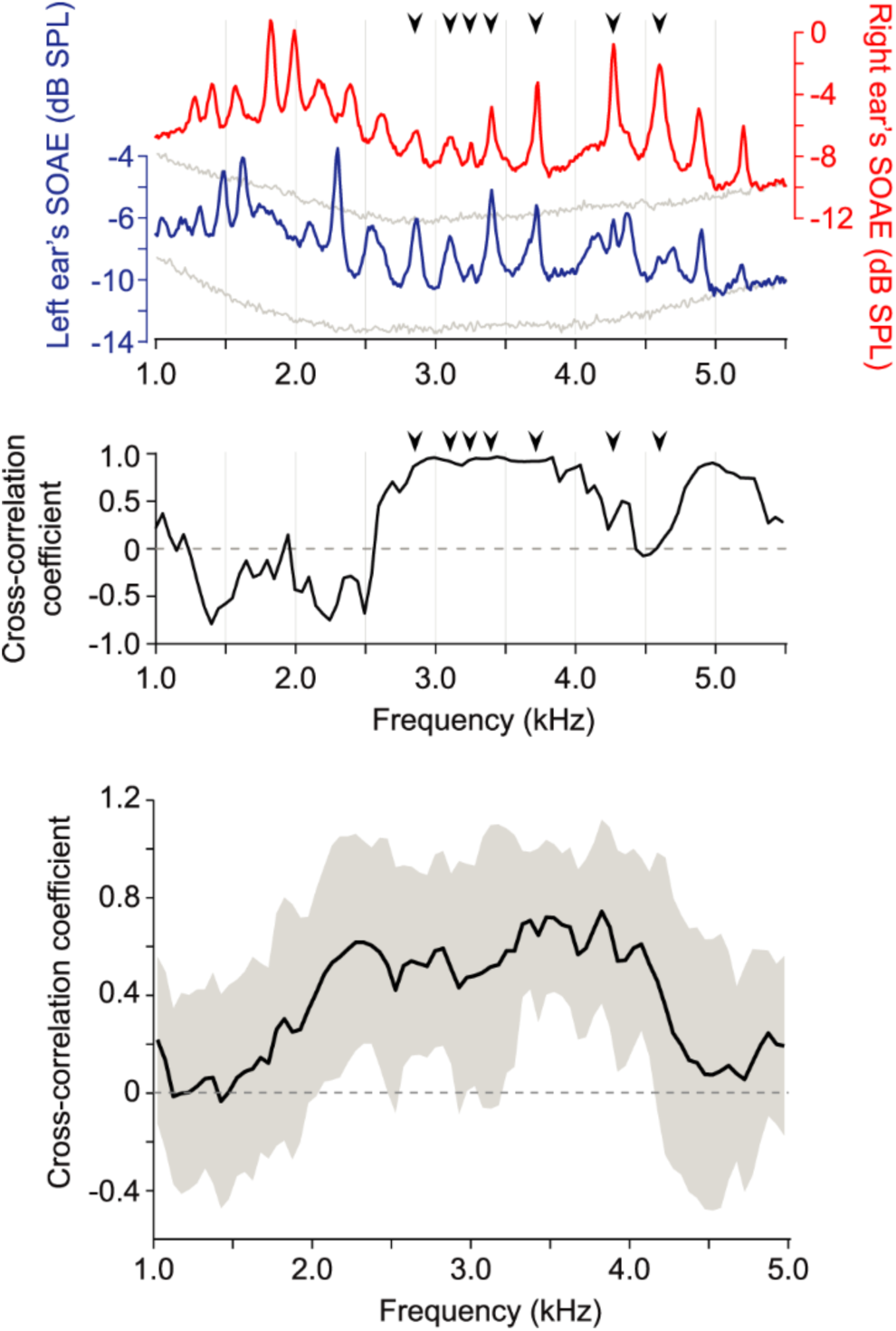
Cross-correlation coefficients between binaural SOAE spectra. (A) Emission spectra recorded simultaneously from both ears of a gecko show identical-frequency SOAEs (black arrowheads). (B) The cross-correlation coefficient is high primarily at frequencies corresponding to identical-frequency SOAEs. Exceptions include those exhibiting a frequency difference from their neighboring monoaural peaks significantly smaller than the width of our analysis window (asterisks). (C) An analysis from 20 geckos reveals that the average cross-correlation coefficients are high between 2 kHz and 4.2 kHz. The shaded area represents standard deviations.

**Figure S4.**
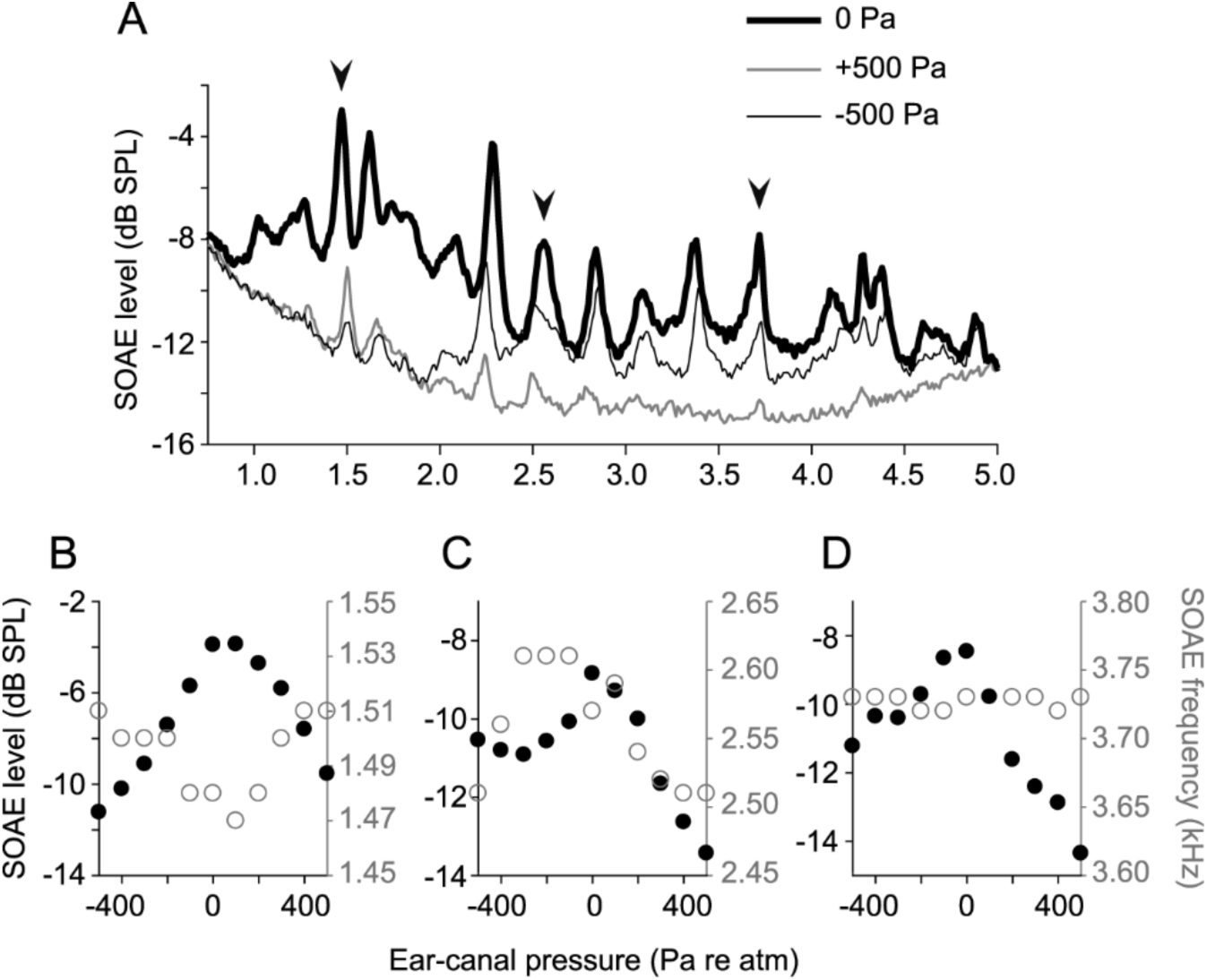
Effects of the ear canal’s static pressure on SOAE level and frequency. (A) SOAEs recorded when the ear-canal gauge pressure was -500 Pa (thin black line) and +500 Pa (thin gray line) were significantly attenuated with respect to those measured at atmospheric pressure (thick black line). (B) Alterations in level and frequency of a low-frequency SOAE peak are symmetric with respect to the sign of the ear-canal pressure change. (C) An SOAE peak in the mid-frequency range was suppressed and shifted by an increase in the ear-canal pressure. (D) At a higher frequency, the level of an SOAE is most sensitive to positive ear-canal gauge pressures. The emission frequency remained unaffected by pressure changes. Black arrowheads in panel A indicate the three SOAE peaks illustrated in panels B-D.

**Figure S5.**
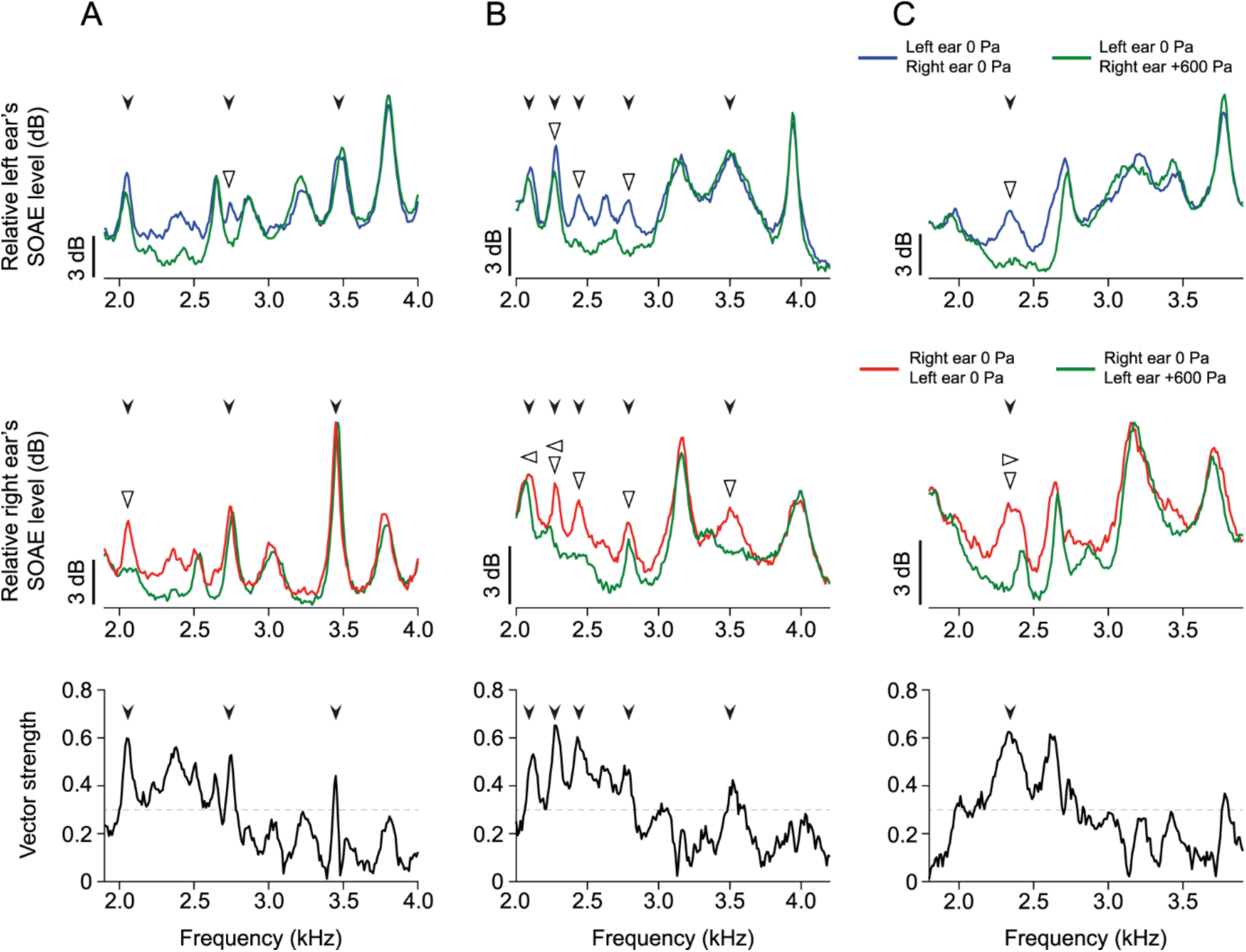
Three examples of alterations in SOAE spectra upon contralateral suppression. (A)-(C) Top panels show the left ear’s SOAE spectra recorded as the right ear-canal pressure was at 0 Pa (blue line) and +600 Pa (green line). Middle panels illustrate the right ear’s SOAE spectra recorded as the left ear-canal pressure was at 0 Pa (red line) and +600 Pa (green line). Black arrowheads indicate identical-frequency SOAEs whose vector strength exceeds 0.3. Changes in the emission level are identified by upright and inverted open triangles and frequency shifts are marked by sideways open triangles. Bottom panels show the vector strengths calculated prior to manipulation of the ear-canal pressure.

### Calculation of *p*-value

A finite-time Fourier transform was performed on each 100 ms nonoverlapping window of the pressure signals recorded before and after manipulations of the contralateral ear-canal pressure. At each frequency component, a student’s t-test was used to compare the spectra. Spectral alterations from contralateral suppression typically corresponded to a *p*-value below 0.001.

**Figure S6.**
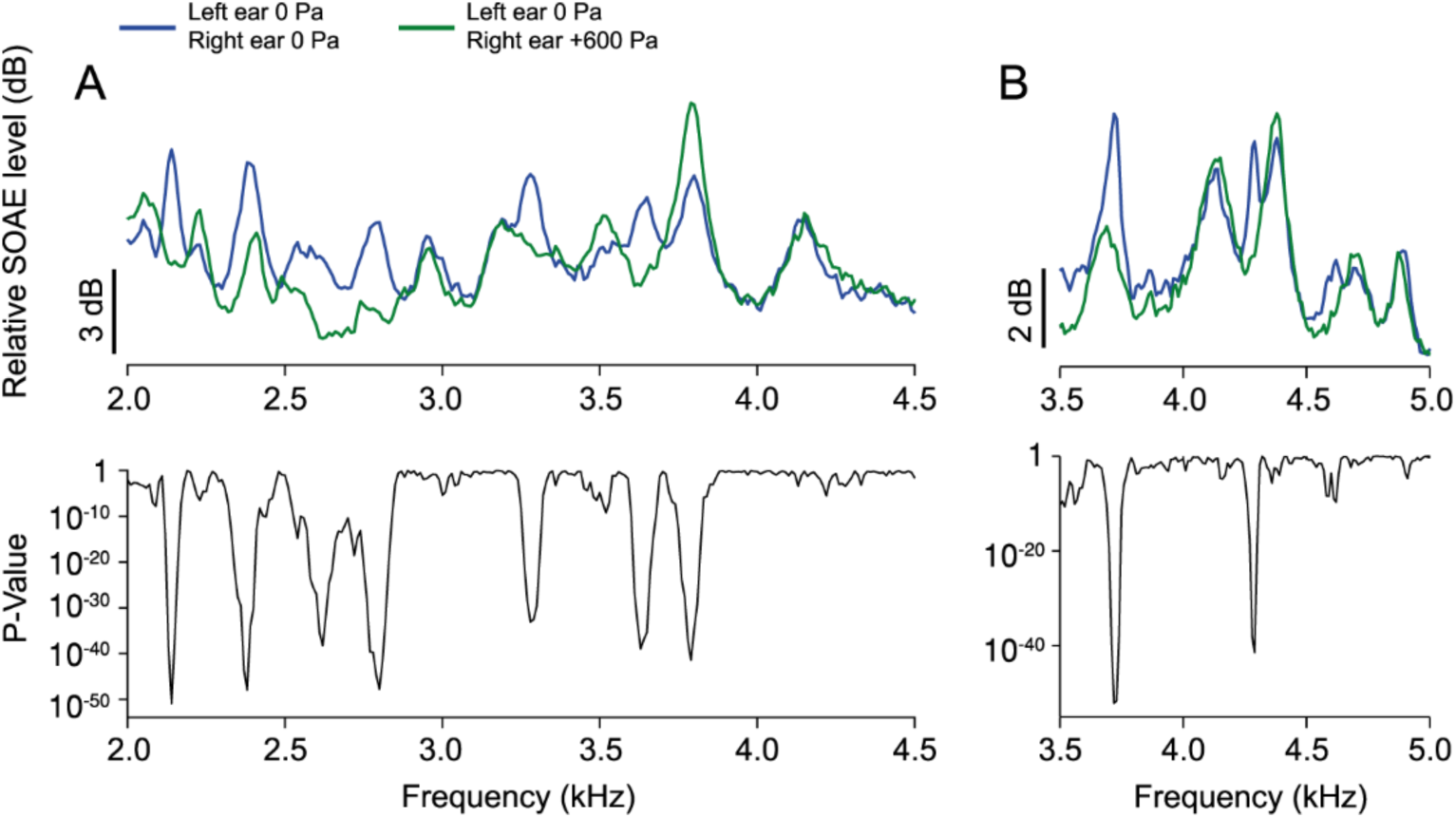
A statistical measure of spectral changes. (A)-(B) Top panels show the left ear’s SOAEs recorded when the right ear-canal gauge pressure was maintained at 0 Pa (blue lines) and +600 Pa (green lines). Bottom panels illustrate the *p*-value for a point-by-point *t*-test between the two emission spectra. Small *p*-values at frequencies corresponding to spectral alterations indicate that the changes in emission levels are statistically significant.

**Figure S7.**
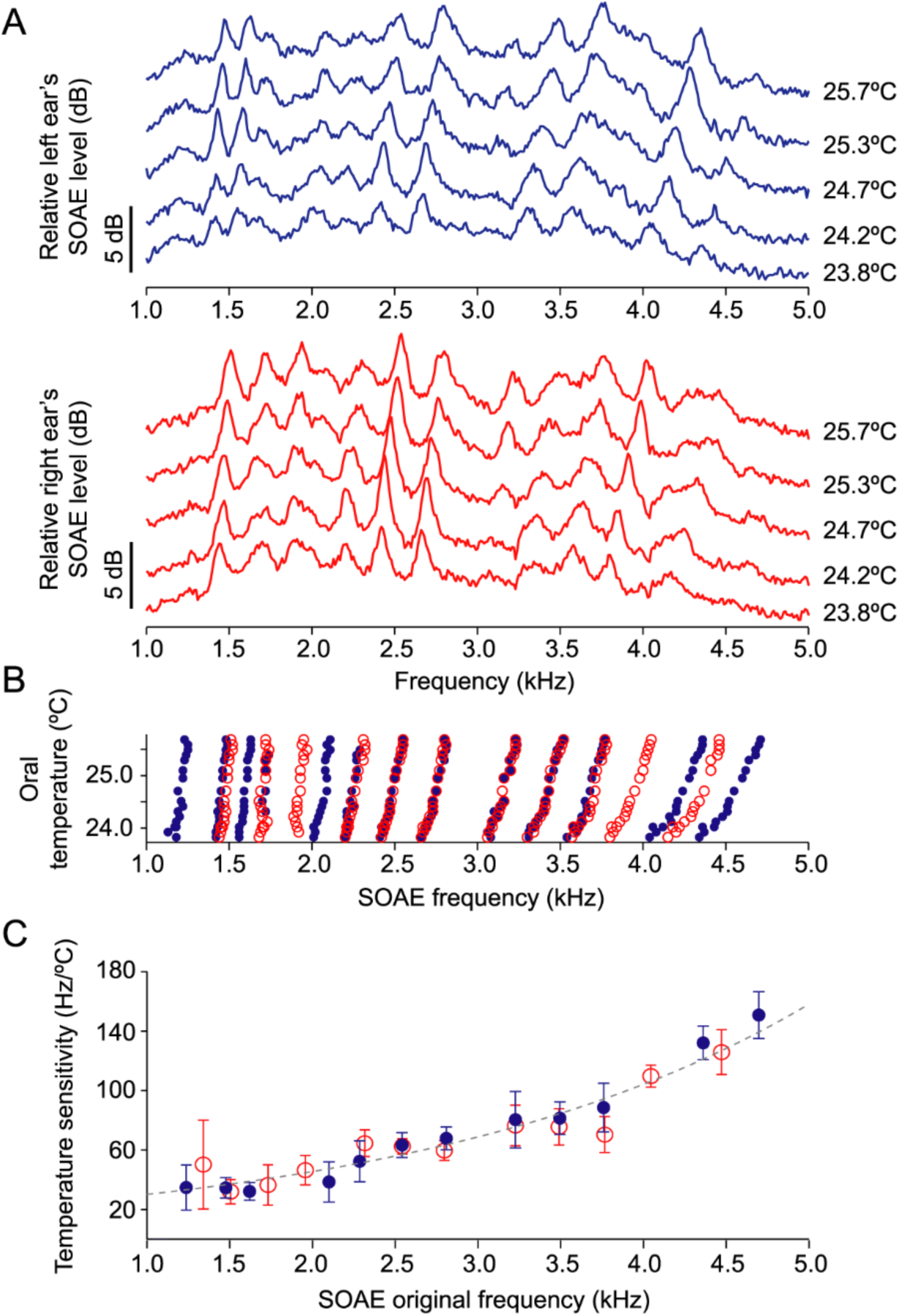
Effects of body temperature on SOAE frequency. (A) SOAEs recorded from the left ear (blue lines) and the right ear (red lines) declined in frequency as the body temperature decreased. The oral temperature was reduced from 25.7 °C to 23.8 °C. (B) Center frequencies of all SOAE peaks from the left ear (blue dots) and the right ear (red circles) systematically shifted toward lower frequencies. (C) The temperature sensitivity of each SOAE peak is the slope of a linear fit to the peak frequency as a function of the oral temperature. The magnitude of the temperature sensitivity is plotted as a function of original SOAE frequency at 25.7 °C (*f*_0_). The shift of SOAEs from both ears can be empirically described by the same exponential function: *df*/*dt* = 18*e*^*f*_0_/1904.8^ (gray dashed line). Error bars indicate the 95 % confidence intervals of the linear fit.

**Figure S8.**
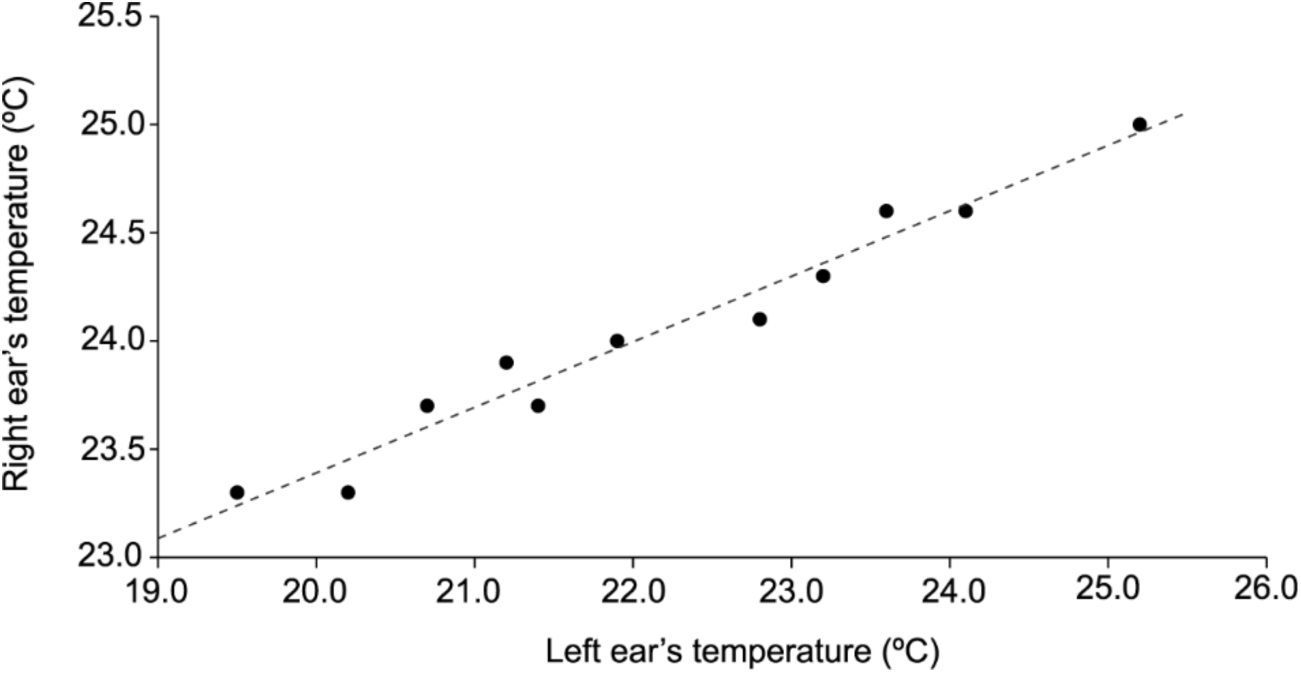
Oral temperatures during asymmetric thermal manipulations. As the left ear was cooled, the right ear’s temperature decreased due to thermal conduction, but always remained higher than the left ear’s temperature. The temperature measured in the oral cavity near the right ear was linearly dependent on that recorded near the left ear: *T*_*right*_ = 0.3026*T*_*left*_ + 17.34.

**Figure S9.**
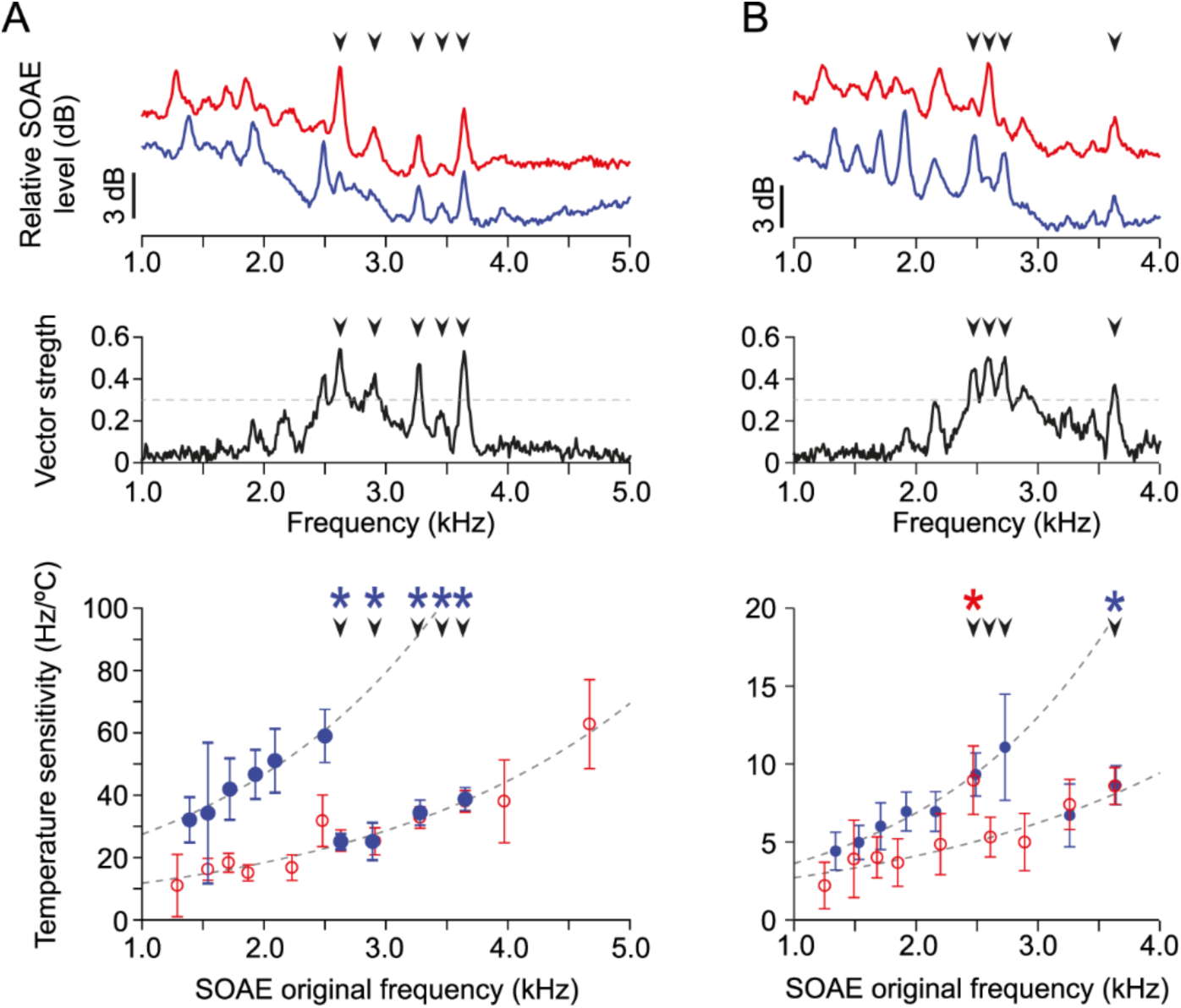
The temperature sensitivity of binaural SOAEs illustrates asymmetric thermal manipulation. (A)-(B) The top panels illustrate SOAE spectra from the left ear (blue lines), and the right ear (red lines). The middle panels display the vector strength before temperature manipulation. The bottom panels show the temperature sensitivity of different SOAE peaks to changes in the left ear’s temperature. After exclusion of identical-frequency SOAEs with high vector strength, the intrinsic temperature sensitivities of each ear could be described by the exponential functions shown in gray dashed lines. Error bars indicate the 95 % confidence intervals of the linear fits. The temperature sensitivity of some highly phase-locked identical-frequency SOAE peaks in the left ear (blue asterisks) equaled that of their counterparts in the right ear. Similarly, a peak in the right ear (red asterisk) possessed a temperature sensitivity that matched that of the left ear. In all panels, black arrowheads indicate highly synchronized identical-frequency SOAEs whose vector strength exceeds 0.3 before temperature manipulation. In (B), a pair of identical-frequency SOAE peaks were originally observed at 2.6 kHz, but the emission from the left ear became undetectable as the temperature was manipulated. Similarly, the right-ear emission from a pair of identical-frequency SOAEs at 2.7 kHz vanished upon asymmetric cooling. Because temperature changes eliminated the contralateral counterparts of these initially synchronized emissions (black arrowheads, no asterisks), their sensitivities matched the intrinsic temperature sensitivity of each ear.

### Coupling noisy oscillators with a damped wave equation

We model the internal coupling of a tokay gecko’s ears by air in its mouth cavity and Eustachian tubes. The cavity is approximated to be a closed cylinder of length *L*, radius *r*, and cross-sectional area *A*^1^. For simplicity, the motion of the air is described in one dimension by its axial velocity *u*(*x, t*) and pressure *p*(*x, t*) with respect to a reference state of zero velocity, pressure *p_r_* = 101 kPa, and temperature *T* = 25°C. To account for damping associated with the air interacting with the sides of the cylinder we respectively write the conservation of momentum and conservation of mass equations as

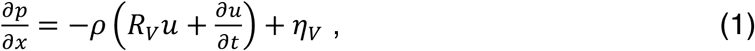

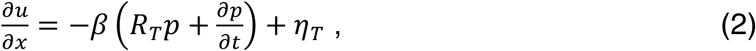

in which *ρ* is the air’s reference density and *β* is the air’s reference adiabatic compressibility. The viscous damping rate is described by

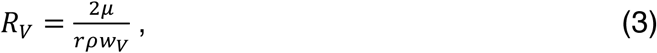

in which *μ* is the dynamic viscosity of the air. The viscous boundary layer width is given by

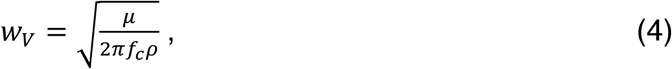

in which *f*_c_ is a characteristic frequency^2^. Similarly, the thermal damping rate is described by

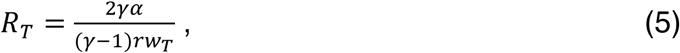

in which *α* is the thermal diffusion rate, *γ* is the air’s adiabatic index, and the thermal boundary layer width is

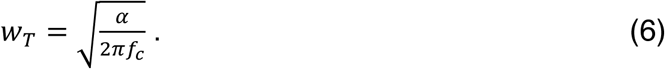

Because the boundary layer widths change slowly with frequency, we fix the characteristic frequency *f*_*c*_ = 15 kHz. To a first approximation, overestimating the characteristic frequency accounts for an increase in the damping rates owing to the intricate anatomy of the head cavity.

According to the fluctuation-dissipation theorem, the Gaussian white noise terms *η*_*v*_(*x, t*) and *η*_*T*_(*x, t*) satisfy

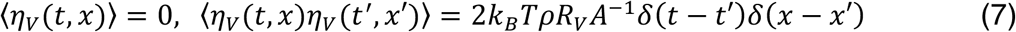

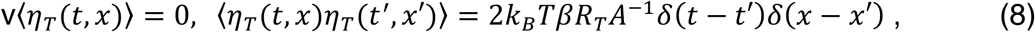

in which 〈 … 〉 denotes the ensemble average and *k*_*B*_ is Boltzmann’s constant^3–4^.

Equations 1 and 2 are noisy telegraph equations, which have been derived previously in the frequency domain^5^. When the noise terms are omitted, the two equations can be combined to yield the acoustic telegraph equation

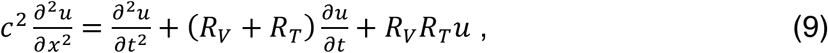

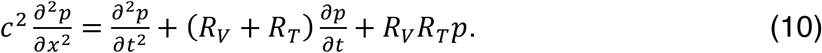

in which 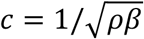 is the reference sound speed^6^.

We used a mathematical description of active hair bundle dynamics to account for the generation of SOAEs within an inner ear^7^. Each eardrum is modeled as an active nonlinear oscillator

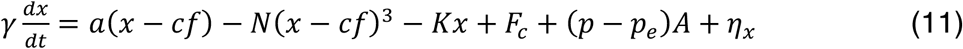

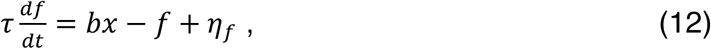

in which *x* denotes the eardrum’s displacement outward from the head cavity. The first three terms on the right-hand side of Equation 11 represent the dynamics of active hair-bundle motility. *a*, *b*, and *K* are stiffnesses, *c* is a compliance, and *N* is the strength of the nonlinearity. *f* is the force due to the active process whose dynamics are described by Equation 12. The oscillator is driven by the difference between the applied pressure in the ear-canal (*p*_*e*_) and the internal pressure (*p*) at the location of the eardrum: *p*(*x* = 0) for the left eardrum and *p*(*x* = *L*) for the right eardrum (Figure 1D). Both pressures are relative to the atmospheric pressure, so *p*_*e*_ = 0 Pa when no additional pressure is applied in the ear canal. *F*_*c*_ is a constant force applied to the oscillator.

The damping coefficient *γ* and time constant *τ* are related to the Gaussian white noise terms *η*_*x*_ and *η*_*f*_ which satisfy

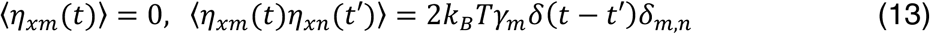

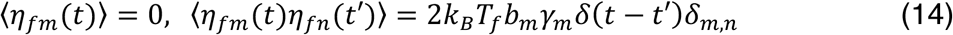

For an active system, the effective temperature *T*_*f*_ may differ from the temperature *T* but was chosen to be equal to *T* here for simplicity.

We couple two active oscillators with displacements *x*_1_ at *x* = 0 for the left eardrum and *x*_2_ at *x* = *L* for the right eardrum (Figure 1D) with the noisy telegraph equations. The boundary condition requires the particle velocities at both ends of the cylinder to be equal to those of the eardrums:

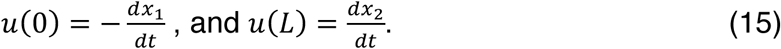

SOAEs are usually measured as external pressure variations in the ear canal. For simplicity, we only describe the displacements of the eardrums and do not calculate the resulting SOAE pressures. In the model, the external pressures in the ear canal (*p*_*e*_) are applied pressures and are not SOAE pressures. SOAE levels are computed from the displacement of the oscillator with respect to the reference level of 3 pm, a displacement chosen to match the velocity of the eardrum at 1 kHz and 0 dB SPL^8–9^. The emission frequency is identical to that of the oscillator.

We solve Equations 1-15 with finite differences and the forward Euler-Maruyama method. To reduce the computation time, we set the noise owing to air fluctuations in Equations 1 and 2 to zero. Excluding this noise is not expected to change the results we present qualitatively. Central differences in time and space were employed for the wave equation and the forward Euler-Maruyama was used for the active oscillators. The space increment was L/100 and the time increment of 100 was reduced to 5 ns when an external static pressure of 600 Pa was applied. Decreasing the increments further had no significant effect on the results. Simulations were performed using C.

The width of a tokay gecko's head^9^ is about 22 mm^2^, but the effective distance between eardrums can be significantly different due to the geometry of the mouth cavity. Based on measurements of the cavity's volume, the fundamental frequency for the cavity of a smaller lizard *Hemidactylus frenatus* has been calculated^1^ to be 5.07 kHz. This calculation assumes the mouth is closed, but the mouth is open in the experiments presented here. Consequently, the fundamental frequency is less than 5.07 kHz and we adjust the effective distance *L* to match the behavior of the phase difference observed in the experiments.

Although the eardrum’s area^8^ is approximately 42 mm^2^, the effective area responsive to acoustic stimulation can be significantly smaller. Because the eardrum is fixed at its perimeter where it connects to the skull, only an area at the center of the membrane oscillates in response to an acoustic stimulation. Therefore, the radius of the tympanum used in this model is about half the actual size.

To change the frequency of an oscillator with little change in its oscillation amplitude, we used a dimensionless timescale parameter *TS* to rescale the oscillator’s time in Equations 1 and 2. Conversely, to change the amplitude without significantly changing the frequency we employed the dimensionless parameter *DS* to rescale the oscillator’s displacement. These changes were achieved by rescaling the oscillators’ standard parameters as *a*_*j*_/*TS*^2^, *b*_*j*_/*TS*^2^, *K*_*j*_/*TS*^2^, *c*_*j*_*TS*^2^, *γ*_*j*_/*TS*, *N*_*j*_/(*TS*^2^*DS*^2^), and *τ*_*j*_*TS*.

To make an oscillator passive we set *c*_*j*_ = 0, the oscillator is no longer coupled to the active force *f*. This passive system was bistable, however, when K = 0.99K_H_ < a. To avoid complications associated with this bistability, we set K = 1.01a in the passive case. Finally, to make the passive oscillator compliant, we set TS = 20.

---

**Table S1.**
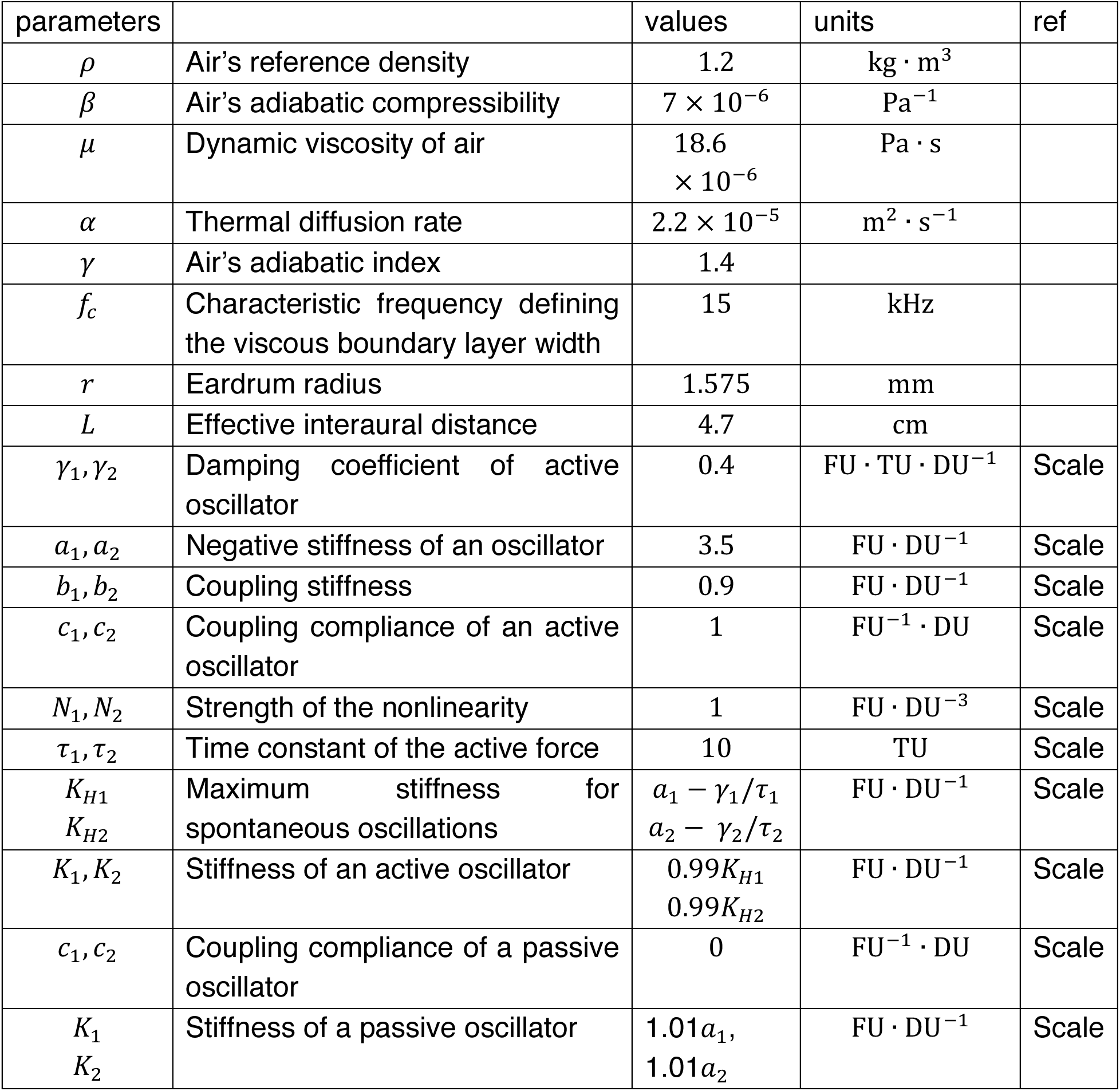
Parameter values used in the model. “Scale” indicates values rescaled from reference, in which the distance unit is DU = 22 pm, the time unit TU = 50 ms, and the force unit FU = 44 nN.

| parameters |  | values | units | ref |
| --- | --- | --- | --- | --- |
| $\rho$ | Air’s reference density | 1.2 | $\text{kg} \cdot \text{m}^3$ | |
| $\beta$ | Air’s adiabatic compressibility | $7 \times 10^{-6}$ | $\text{Pa}^{-1}$ | |
| $\mu$ | Dynamic viscosity of air | $18.6 \times 10^{-6}$ | $\text{Pa} \cdot \text{s}$ | |
| $\alpha$ | Thermal diffusion rate | $2.2 \times 10^{-5}$ | $\text{m}^2 \cdot \text{s}^{-1}$ | |
| $\gamma$ | Air’s adiabatic index | 1.4 | | |
| $f_c$ | Characteristic frequency defining the viscous boundary layer width | 15 | kHz | |
| $r$ | Eardrum radius | 1.575 | mm | |
| $L$ | Effective interaural distance | 4.7 | cm | |
| $\gamma_1, \gamma_2$ | Damping coefficient of active oscillator | 0.4 | $FU \cdot TU \cdot DU^{-1}$ | Scale |
| $a_1, a_2$ | Negative stiffness of an oscillator | 3.5 | $FU \cdot DU^{-1}$ | Scale |
| $b_1, b_2$ | Coupling stiffness | 0.9 | $FU \cdot DU^{-1}$ | Scale |
| $c_1, c_2$ | Coupling compliance of an active oscillator | 1 | $FU^{-1} \cdot DU$ | Scale |
| $N_1, N_2$ | Strength of the nonlinearity | 1 | $FU \cdot DU^{-3}$ | Scale |
| $\tau_1, \tau_2$ | Time constant of the active force | 10 | TU | Scale |
| $K_{H1}, K_{H2}$ | Maximum stiffness for spontaneous oscillations | $a_1 - \gamma_1/\tau_1$<br>$a_2 - \gamma_2/\tau_2$ | $FU \cdot DU^{-1}$ | Scale |
| $K_1, K_2$ | Stiffness of an active oscillator | $0.99K_{H1}$<br>$0.99K_{H2}$ | $FU \cdot DU^{-1}$ | Scale |
| $c_1, c_2$ | Coupling compliance of a passive oscillator | 0 | $FU^{-1} \cdot DU$ | Scale |
| $K_1, K_2$ | Stiffness of a passive oscillator | $1.01a_1,$<br>$1.01a_2$ | $FU \cdot DU^{-1}$ | Scale |

## References

1 Ohyama, K., Wada, H., Kobayashi, T. & Takasaka T. Spontaneous otoacoustic emissions in the guinea pig. Hear. Res. 56, 111–121 (1991).

2 Probst, R., Lonsbury-Martin, B. L. & Martin, G. K. A review of otoacoustic emissions. J. Acoust. Soc. Am. 89, 2027–2067 (1991).

3 van Dijk, P., Narins, P. M. & Wang, J. Spontaneous otoacoustic emissions in seven frog species. Hear. Res. 101, 102–112 (1996).

4 Manley, G. A., Gallo, L. & Koppl, C. Spontaneous otoacoustic emissions in two gecko species, Gekko gecko and Eublepharis macularius. J. Acoust. Soc. Am. 99, 1588–1603 (1996).

5 Taschenberger, G. & Manley, G. A. Spontaneous otoacoustic emissions in the barn owl. Hear. Res. 110, 61–76 (1997).

6 Manley, G. A. Evidence for an active process and a cochlear amplifier in nonmammals. J. Neurophysiol. 86, 541–549 (2001).

7 Koppl, C. & Manley, G. A. Spontaneous otoacoustic emissions in the bobtail lizard. II: Interactions with external tones. Hear. Res. 72, 159–170 (1994).

8 Long, G. R., Tubis, A. & Jones, K. L. Modeling synchronization and suppression of spontaneous otoacoustic emissions using Van der Pol oscillators: effects of aspirin administration. J. Acoust. Soc. Am. 89, 1201–1212 (1991).

9 Long, G. R., van Dijk, P. & Wit, H. P. Temperature dependence of spontaneous otoacoustic emissions in the edible frog (Rana esculenta). Hear. Res. 98, 22–28 (1996).

10 Hauser, R., Probst, R. & Harris, F. P. Effects of atmospheric pressure variation on spontaneous, transiently evoked, and distortion product otoacoustic emissions in normal human ears. Hear. Res. 69, 133–145 (1993).

11 van Dijk, P., Maat, B. & de Kleine, E. The effects of static ear canal pressure on human spontaneous otoacoustic emissions: Spectral width as a measure of the intracochlear oscillation amplitude. J. Assoc. Res. Otolaryngol. 12, 13–28 (2011).

12 van Dijk, P. & Manley, G. A. The Effects of Air Pressure on Spontaneous Otoacoustic Emissions of Lizards. J. Assoc. Res. Otolaryngol. 14, 309–319 (2013).

13 Ó Maoiléidigh, D., Nicola, E. M. & Hudspeth, A. J. The diverse effects of mechanical loading on active hair bundles. Proc. Natl. Acad. Sci. 109, 1943–1948 (2012).

14 Penner, M. J., Brauth, S. E. & Jastreboff, P. J. Covariation of binaural, concurrently-measured spontaneous otoacoustic emissions. Hear. Res. 73, 190–194 (1994).

15 Braun, M. Accurate binaural mirroring of spontaneous otoacoustic emissions suggests influence of time-locking in medial efferents. Hear. Res. 118, 129–138 (1998).

16 van Dijk, P., Wit, H. P. & Segenhout, J. M. Spontaneous otoacoustic emissions in the European edible frog (Rana esculenta): Spectral details and temperature dependence. Hear. Res. 42, 273–282 (1989).

17 Christensen-Dalsgaard, J. & Manley, G. A. Directionality of the lizard ear. J. Exp. Biol. 208, 1209–1217 (2005).

18 Christensen-Dalsgaard, J. & Manley, G. A. Acoustical coupling of lizard eardrums. J. Assoc. Res. Otolaryngol. 9, 407–416 (2008).

19 Christensen-Dalsgaard, J., Tang, Y. & Carr, C. E. Binaural processing by the gecko auditory periphery. J. Neurophysiol. 105, 1992–2004 (2011).

20 Martin, P. & Hudspeth A. J. Active hair-bundle movements can amplify a hair cell’s response to oscillatory mechanical stimuli. Proc. Natl. Acad. Sci. 96, 14306–11 (1999).

21 Kennedy, H. J., Crawford, A. C. & Fettiplace, R. Force generation by mammalian hair bundles supports a role in cochlear amplification. Nature 433, 880–883 (2005).

22 Nadrowski, B., Martin P. & Jülicher F. Active hair-bundle motility harnesses noise to operate near an optimum of mechanosensitivity. Proc. Natl. Acad. Sci. 101, 12195–12200 (2004).

23 Fredrickson-Hemsing, L., Ji S., Bruinsma, R. & Bozovic, D. Mode-locking dynamics of hair cells of the inner ear. Phys Rev E. 86:021915 (2012).

24 Salvi, J. D., Ó Maoiléidigh, D., Fabella B. A., Tobin, M. & Hudspeth, A. J. Control of a hair bundle’s mechanosensory function by its mechanical load. Proc. Natl. Acad. Sci. 112, E1000–E1009 (2015).

25 Martin, P. et al. Spontaneous oscillation by hair bundles of the bullfrog’s sacculus. J. Neurosci. 23, 4533–48 (2003).

26 Till, B. C. & Driessen P. F. A didactically novel derivation of the telegraph equation to describe sound propagation in rigid tubes. Eur. J. Phys. 35, 015007 (2014).

27 Köppl, C. & Manley, G. A. Spontaneous otoacoustic emissions in the bobtail lizard. III: Temperature effects. Hear. Res. 72, 171–180 (1994).

28 Wever, E. G. The reptile ear-Its structure and function (Princeton Univ. Press, Princeton, 1978).

29 Salvi, J. D., Ó Maoiléidigh, D. & Hudspeth A. J. Identificantion of bifurcations from observations of noisy biological oscillators. Biophys. J. 111, 798–812 (2016).

30 Vossen, C., Christensen-Dalsgaard, J. & van Hemmen, J. L. Analytical model of internally coupled ears. J. Acoust. Soc. Am. 128, 909–918 (2010).

31 Manley, G. A. The middle ear of the tokay gecko. J. Comp. Physiol. 81, 239–250 (1972).

32 Miller M. R. A scanning electron microscope study of the papilla basilaris of Gekko gecko. Z. Zellforsch. 136, 307–328 (1973)

33 Manley G. A., Köppl C. & Sneary M. Reversed tonotopic map of the basilar papilla in Gekko gecko. Hear. Res. 131, 107–116 (1999).

34 Köppl C. & Authier S. Quantitative anatomical basis for a model of micromechanical frequency tuning in the Tokay gecko, Gekko gecko. Hear. Res. 82, 14–25 (1995).

## References

1 Vossen, C., Christensen-Dalsgaard, J. & van Hemmen, J. L. Analytical model of internally coupled ears. J. Acoust. Soc. Am. 128, 909–918 (2010).

2 Keefe, D. H. Acoustical wave propagation in cylindrical ducts: Transmission line parameter approximations for isothermal and nonisothermal boundary conditions. J. Acoust. Soc. Am. 75, 58–62 (1984).

3 Nyquist H. Thermal agitation of electric charge in conductors. Phys. Rev. 32, 110–113 (1928).

4 Callen, H. B. & Welton, T. A. Irreversibility and generalized noise. Phys. Rev. 83, 34–40 (1951).

5 Zurita-Sanchez, J. R. & Henkel, C. Lossy electrical transmission lines: thermal fluctuations and quantization. Phys. Rev. A. 73, 063825 (2006).

6 Till, B. C. & Driessen, P. F. A didactically novel derivation of the telegraph equation to describe sound propagation in rigid tubes. Eur. J. Phys. 35 015007 (2014).

7 Ó Maoiléidigh, D., Nicola, E. M. & Hudspeth, A. J. The diverse effects of mechanical loading on active hair bundles. Proc. Natl. Acad. Sci. 109, 1943–1948 (2012).

8 Manley, G. A. The middle ear of the tokay gecko. J. Comp. Physiol. 81, 239–250 (1972).

9 Christensen-Dalsgaard, J. & Manley, G. A. Directionality of the lizard ear. J. Exp. Biol. 208, 1209–1217 (2005).

